# Highly polygenic control of photosynthetic responses to nighttime temperature in Arabidopsis studied by genomic prediction

**DOI:** 10.1101/2025.11.08.687332

**Authors:** Ana Carolina dos Santos Sá, Alina Bazarova, Andreas Fischbach, Stefan Kesselheim, Benjamin Stich, Shizue Matsubara

## Abstract

- Rising nighttime temperature (T_night_) can reduce crop yields, while low T_night_ may restrict plant growth and development. Despite these quantifiable effects of T_night_, the genetic basis underlying plant responses to T_night_ remains unclear. We investigated natural variation in long-term response of effective photosynthetic efficiency (F_q_’/F_m_’) to T_night_ among Arabidopsis accessions.
- Genome-wide association study (GWAS) was conducted for F_q_’/F_m_’ of the accessions grown under 15°C or 20°C T_night_. The associated single nucleotide polymorphisms (SNPs) were identified and incorporated in genomic prediction (GP) models to assess the improvement of prediction accuracy. The predictions were experimentally validated in an independent, genetically diverse population.
- GWAS revealed highly polygenic architecture of F_q_’/F_m_’, with associated SNPs varying across T_night_ conditions and measurement days. Notably, 15°C T_night_ stabilized the contributions of a subset of associated SNPs, whereas 20°C T_night_ enhanced day-to-day variations in SNP-trait associations. The GWAS-derived SNPs significantly improved the prediction accuracy of GP models, indicating their collective influence. The validation experiment confirmed the identification of low-F_q_’/F_m_’ accessions in 15°C T_night_.
- The results uncover the genetic underpinnings of long-term F_q_’/F_m_’ response to cool *vs* warm nights and establish a framework for leveraging GWAS and GP to explore complex traits, such as photosynthesis, toward breeding climate-resilient crops.

## Introduction

Global climate change is increasing nighttime temperature (T_night_) more rapidly than daytime temperature (T_day_)^1–3^. Episodes of elevated T_night_ can cover larger geographic areas and last longer than those of high T_day_^4^. While heatwaves and maximal T_day_ have gained widespread attention in plant science and crop research, the impacts of nighttime warming are understudied. Yet, high T_night_ can reduce yields of staple crops such as wheat^5^, rice^6,7^, and barley^5^ and pose a threat to agricultural sustainability and food security.

The mechanisms underlying yield losses at elevated T_night_ are complex, including altered carbon allocation due to increased nocturnal respiration^7^ and decreased sugar export from leaves^8^, as well as a shorter grain-filling period^5,9^. Besides the direct effects of T_night_ on nocturnal carbon partitioning and utilization, previous studies have highlighted (indirect) effects on daytime photosynthesis^8,10,11^. Whilst high T_night_ may decrease crop yields by curtailing resource allocation to reproductive organs, low T_night_ can limit plant growth and biomass accumulation^11^. Growth suppression at low T_night_ is often accompanied by a decline in photosynthesis, even at moderately low T_night_^4,8,12^. Despite these quantifiable effects of T_night_ on growth, photosynthesis, and yields, the genetic underpinnings of plant responses to T_night_ remain poorly understood.

The analysis of natural genetic variation offers valuable insights into genetic architecture underlying a range of plant traits^13^. Genome-wide association study (GWAS) ‒an approach that statistically tests for associations between genetic markers and phenotypic variation^14^‒ uncovers trait-associated genetic variants and therewith functional polymorphisms. Although GWAS has proven powerful in identifying allelic variants associated with important agricultural traits, unravelling the genetic architecture of polygenic traits, which are governed by numerous small-effect variants, remains a major challenge^14^. Meanwhile, a paradigm shift has occurred in the study of complex traits, from identifying genetic variants with strong statistical associations to developing accurate prediction models^15^.

Genomic prediction (GP) builds a model using genome-wide markers and phenotypic data to estimate breeding values for genotypes with marker information alone^16^. Genomic best linear unbiased prediction (gBLUP) has been widely used in plant and animal breeding^17,18^; however, because of its additive nature, alternative approaches have been developed to capture complex, non-additive interactions among markers, such as epistasis^19,20^. Examples of non-linear models include ensemble machine learning methods which combine multiple weak models into a single, more robust and accurate model^21^. Whether non-linear models can consistently surpass linear models in predicting complex polygenic traits remains an unsettled question in quantitative genetics^19,20^.

Here, we explored natural genetic variation underlying long-term photosynthetic responses to two T_night_ conditions (15°C and 20°C) in *Arabidopsis thaliana*. Our GWAS revealed numerous and distinct single nucleotide polymorphisms (SNPs) associated with variations in effective photosystem II (PSII) efficiency (F_q_’/F_m_’) among the accessions grown under 15°C and 20°C T_night_. Midday F_q_’/F_m_’ showed a highly polygenic architecture, with hundreds of small-effect SNPs contributing variably across different days and T_night_ conditions. Incorporating these GWAS-derived SNPs into linear and non-linear GP models markedly improved prediction accuracy in both T_night_ conditions, highlighting their collective influence on F_q_’/F_m_’. Remarkably, 15°C T_night_ stabilized the contributions of a subset of associated SNPs, whereas 20°C T_night_ induced larger day-to-day variation in SNP-trait associations. A validation experiment confirmed the models’ capacity to identify low-F_q_’/F_m_’ accessions in an independent, genetically diverse population, demonstrating the potential of this approach for advancing our understanding and prediction of complex traits.

## Results

### Natural genetic variation in long-term response of photosynthesis to T_night_

To assess genetic diversity in long-term photosynthetic response to T_night_, we measured PSII efficiency in 308 Arabidopsis accessions (Supplementary Table 1) from the HapMap population^22^ grown in 15°C and 20°C T_night_ condition. Both T_night_ treatments had the same T_day_ of 26°C.

The adjusted entry-means of F_q_’/F_m_’ measured around midday under the maximal growth light intensity of ca. 600 μmol photons m^-2^ s^-1^ were, on average across all accessions, higher in the plants grown in 20°C than 15°C T_night_ (Fig. 1A). The opposite was the case for the maximal PSII efficiency (F_v_/F_m_) measured during the night at the respective T_night_, with slightly higher values found in 15°C T_night_ (Fig. 1B). The accessions showed large phenotypic variance for PSII efficiency in both T_night_ conditions, with larger variations for light-adapted midday F_q_’/F_m_’ than for dark-adapted F_v_/F_m_ (Fig. 1A, B; Supplementary Table 2). Broad-sense heritability (defined as the proportion of genetic variance to phenotypic variance) of the PSII efficiency parameters was generally high in our experimental conditions (Fig. 1E), especially for F_q_’/F_m_’. The heritability of these parameters was higher in 20°C than in 15°C T_night_ and increased in both conditions from 16 to 18 days after stratification (DAS) (Fig. 1E).

**Figure 1.**
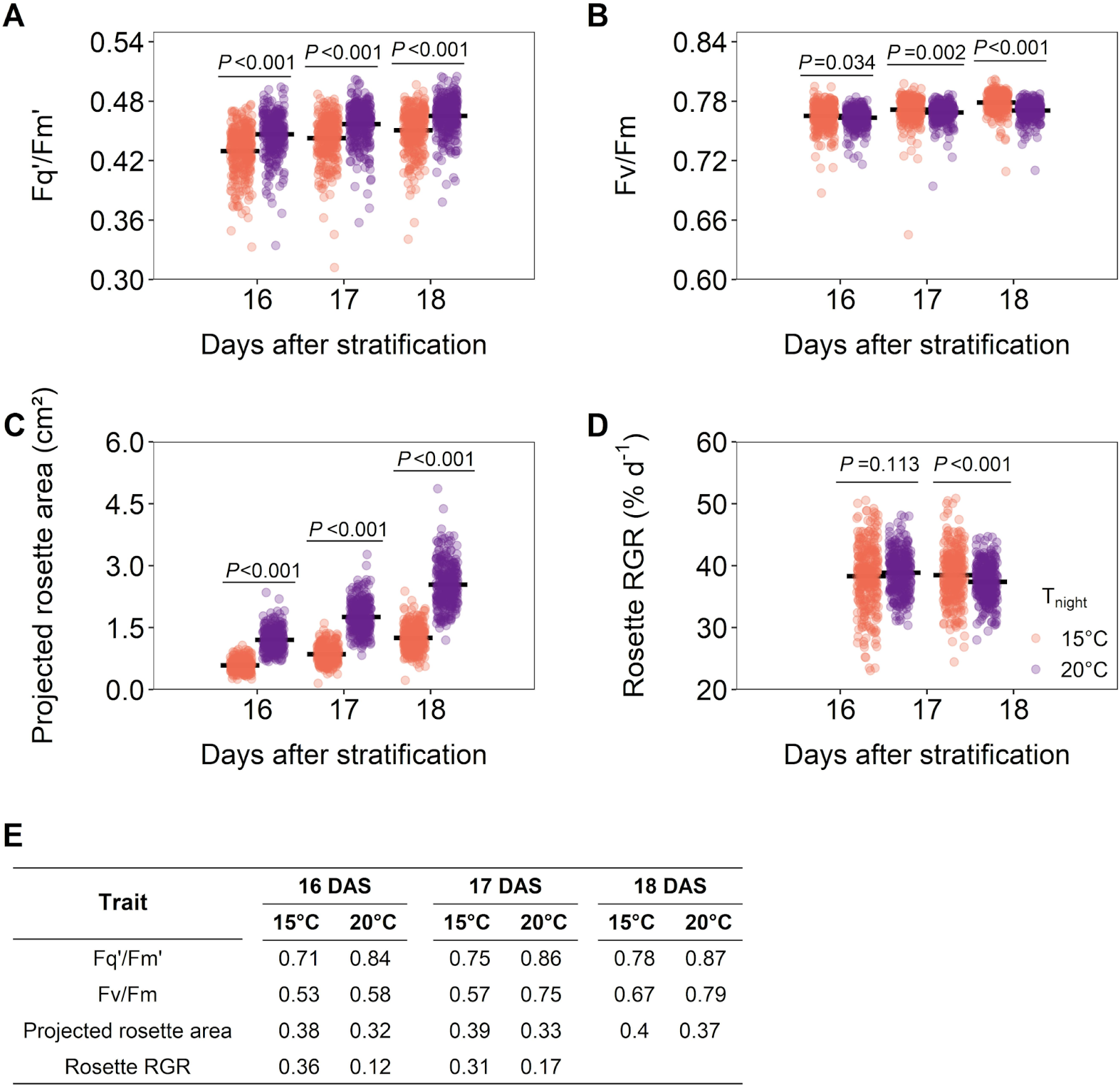
Effects of night temperature (T_night_) on PSII efficiency and growth traits. A and B, the effective (F_q_’/F_m_’) and the maximal (F_v_/F_m_) PSII efficiency. C and D, projected rosette area and relative growth rate (RGR). E, broad-sense heritability of these traits. Traits were measured in 308 Arabidopsis accessions from the HapMap population (listed in Supplementary Table 1) grown in the 15°C (coral) and 20°C (purple) T_night_ conditions at 16, 17 and 18 days after stratification (DAS). F_q_’/F_m_’ was measured at around midday under the maximal growth light intensity of ∼600 μmol photons m^-2^ s^-1^ and daytime growth temperature of 26°C, while F_v_/F_m_ was measured at respective T_night_ during the night. The rosette RGR was calculated from the projected rosette area measured on two consecutive days (i.e., RGR at 16 DAS was calculated from the projected rosette area at 16 and 17 DAS). Horizontal lines in panel A-D show means of all accessions. Circles represent adjusted entry-means of individual accessions (n=2-10 for each accession). Significant differences between the two treatments were tested by two-tailed t-test.

Temperature has a strong impact on plant growth. As expected, Arabidopsis plants were smaller in 15°C than in 20°C T_night_ (Fig. 1C). In comparison, the effects of T_night_ were less obvious for relative growth rate (RGR) of rosettes (Fig. 1D). While accessions displayed large phenotypic variance also for these growth parameters, their heritability was much lower than that of PSII efficiency (Fig. 1E).

### T_night_ affects genetic architecture of F_q_’/F_m_’

Having seen the large phenotypic variance and high heritability of PSII efficiency in the plants grown in 15°C and 20°C T_night_, we conducted GWAS to map genetic variants underlying the natural variation in midday F_q_’/F_m_’ under these T_night_ conditions. Of the 308 accessions that were phenotyped, 293 were included in GWAS since SNP information was not available for the remaining 15 accessions.

Principal component (PC) analysis was performed for the 293 accessions based on 211,771 SNPs, excluding rare SNPs with an allele frequency below 2%. PC1 and PC2 explained 3.1% and 2%, respectively, of the total genetic variance among these accessions (Supplementary Fig. 1). Up to five PCs were included in the GWAS models as covariates to correct for population structure (see Q-Q plots in Supplementary Fig. 2). The SNP-trait associations identified by the model FarmCPU were considered significant when -log_10_ (*P*) ≥ FDR (false discovery rate) threshold (0.05) and/or -log_10_ (*P*) > 4.0 on at least two days (Supplementary Fig. 3; Supplementary Table 3). No more than 13 SNPs met the latter criterion in 15°C T_night_, of which SNP No. 2 was identified on all three days (Fig. 2). None of these SNPs met the former criterion in 15°C T_night_. In 20°C T_night_, 23 SNPs met the former criterion, of which four (No. 15, 16, 20 and 27) also met the latter and No. 15 was identified on all three days (Fig. 2). Except SNP No. 2 which was detected in both 15°C and 20°C T_night_, the SNP-trait associations did not overlap between the two conditions (Fig. 2).

**Figure 2.**
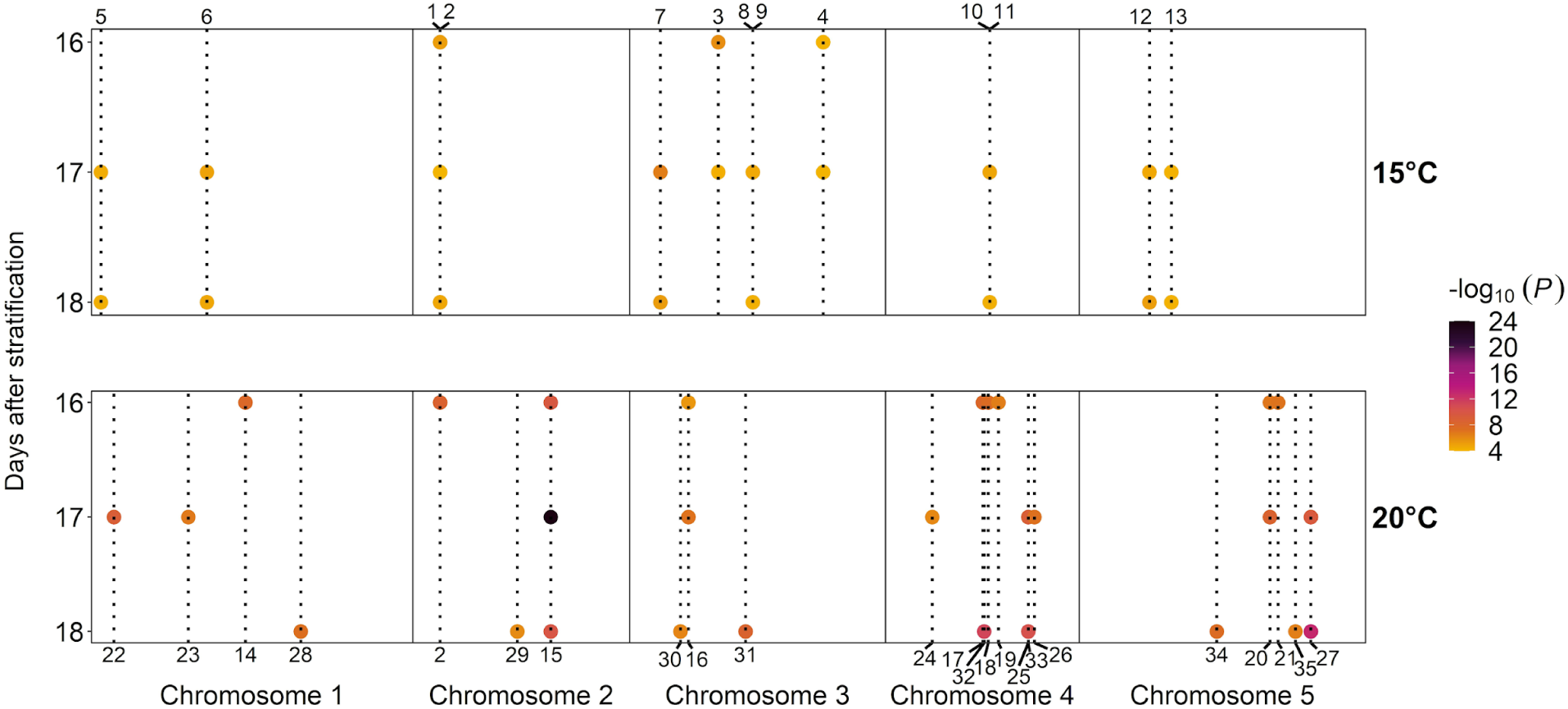
Summary of single nucleotide polymorphisms (SNPs) associated with long-term acclimation of F_q_’/F_m_’ to night temperature. Genome-wide SNP-trait association was analyzed based on F_q_’/F_m_’ measured in the 293 Arabidopsis accessions grown in 15°C and 20°C T_night_ at 16, 17 and 18 days after stratification. SNP-trait associations were considered significant when -log_10_ (*P*) ≥ FDR (false discovery rate) threshold (0.05) and/or -log_10_ (*P*) > 4.0 on at least two dates. Circles represent individual SNPs. Colors of the circles show significance of association (*P*). SNPs are numbered from No. 1 to No. 13 for 15°C T_night_ and from No. 14 to No. 35 for 20°C T_night_. Information on these SNPs is provided in Supplementary Table 3.

The identified SNPs explained together 18-32% of the F_q_’/F_m_’ variance in 15°C T_night_ and 40-46% in 20°C T_night_ on different days (see r^2^ of simultaneous fits in Supplementary Table 3). Contributions of individual SNPs ranged from 5 to 12% in 15°C T_night_ and from 2 to 19% in 20°C T_night_, with the highest r^2^ for SNP No. 7 in 15°C T_night_ and SNP No. 15 in 20°C T_night_ at 17 DAS. These SNPs also had the largest modulus of the effect (given in the unit of F_q_’/F_m_’), -0.022 for No. 7 and -0.024 for No. 15 (Supplementary Table 3). Most of the other SNPs had estimated effect size of ± 0.01 or smaller.

None of the identified SNPs is located in loci known to be involved in photosynthesis (Supplementary Table 3). Of the aforementioned SNPs, for instance, SNP No. 2 is found in AT2G03730 encoding an ACT domain containing protein (ACT DOMAIN REPEAT 5), No. 7 in AT3G05050 encoding a protein kinase superfamily protein, and No. 15 in AT2G26250 encoding an enzyme 3-ketoacyl-CoA synthase (3-KETOACYL-COA SYNTHASE 10, aka FIDDLEHEAD) involved in the biosynthesis of very-long-chain fatty acids in epidermis and phloem^23^.

### GWAS-derived SNP-trait associations improve GP

If the variants identified by GWAS contribute to the genetic variance in F_q_’/F_m_’ among the accessions, information on these SNPs should enhance the power of GP to predict F_q_’/F_m_’ in these genotypes under the treatment conditions. Thus, to corroborate the collective influence of the GWAS-derived SNPs on F_q_’/F_m_’, we evaluated the performance of gBLUP by using three different sets of SNPs as predictors: i) 211,771 SNPs in the Arabidopsis GWAS panel, ii) GWAS-derived SNPs with -log_10_ (*P*) ≥ FDR threshold (0.05) and/or -log_10_ (*P*) > 4.0 on at least two days (Supplementary Table 3), and iii) GWAS-derived SNPs with -log_10_ (*P*) ≥ 3.0 on each day (Supplementary Data). The first set is for prediction without the information on GWAS-derived SNP-trait associations. The third set is based on a liberal significance threshold but contains more information on SNP-trait associations compared to the second set (Supplementary Table 4).

Prediction ability, defined as the median Pearson correlation coefficient between the actual and the predicted F_q_’/F_m_’ across all cross-validation runs, varied substantially depending on the SNP sets used (Fig. 3). Low median prediction ability (∼0.33) was obtained by using the 211,771 SNPs, with large variations across the test sets (Fig. 3A). Since gBLUP assumes equal contributions of individual genetic markers to the phenotype, we tested whether the prediction can be improved by the Bayesian (Bayes) approach that allows for differential distribution of marker effects^16^. The results, however, indicated no improvement of predictive performance by Bayes A and Bayes B compared to gBLUP (Supplementary Table 5). Next, we used the information of the second set in the gBLUP model. This SNP set moderately increased the prediction ability, yielding a median prediction ability of 0.51 in 15°C T_night_ and 0.65 in 20°C T_night_, but the performance was still varying widely among the test sets (Fig. 3B). Integrating the third set, which included a larger number of less strongly associated SNPs compared to the second set, markedly improved the prediction ability to reach above 0.8, while at the same time diminishing the disparity among the test sets in both T_night_ conditions (Fig. 3C).

**Figure 3.**
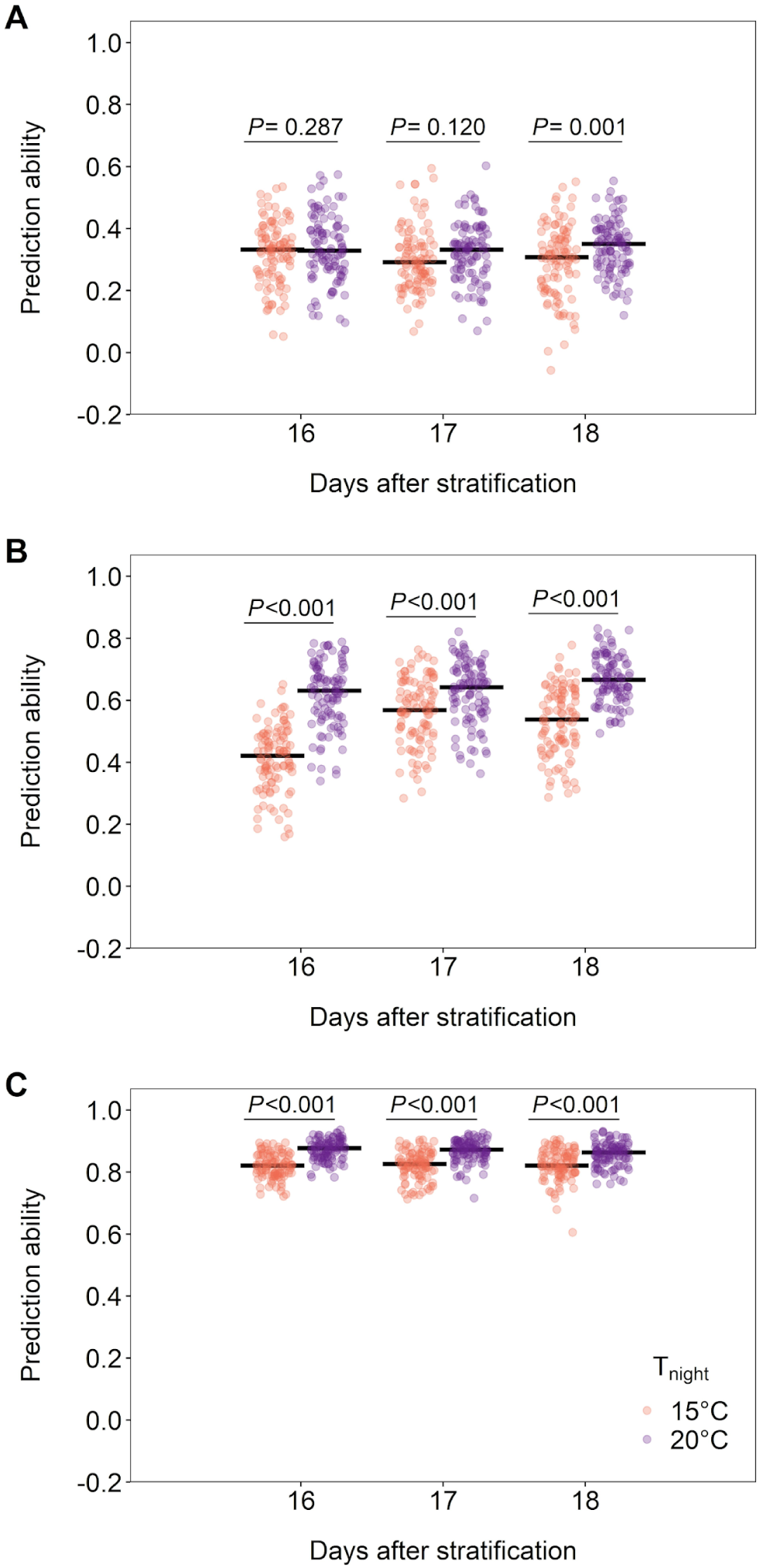
Prediction ability of F_q_’/F_m_’. The F_q_’/F_m_’ data of 293 accessions were used to train gBLUP models based on 211,771 SNPs (A), GWAS-derived SNPs with -log_10_ (*P*) ≥ FDR threshold and/or -log_10_ (*P*) > 4.0 in at least two days (B), or with -log_10_ (*P*) ≥ 3.0. Horizontal bars show the median of 100 cross-validation replications. Circles represent individual replications in 15°C (coral) or 20°C (purple) T_night_. Statistically significant differences in prediction ability between the treatments were tested by two-tailed t-test.

The higher prediction ability by the third set than the first set is attributable to higher SNP-trait associations; predictions based on the same number of randomly selected, weakly associated SNPs (-log_10_ (*P*) < 3.0) showed poor prediction abilities compared to the SNPs in the third set (-log_10_ (*P*) ≥ 3.0) (Supplementary Fig. 4). The higher prediction ability by the third set than the second set, on the other hand, may be due to the tendency of gBLUP to predict the phenotype more accurately with increasing number of SNPs even if each SNP explains only a small proportion of the variance^24,25^. Indeed, when the prediction ability was compared between the second and the third sets based on the same number of SNPs as in the second set (see Supplementary Table 4 for the number of SNPs), randomly chosen SNPs from the third set underperformed in 20°C T_night_ (Supplementary Fig. 5B). It seems that the inclusion of additional SNPs in the third set, despite the moderate significance of their associations, increased the prediction ability in this condition. In contrast, the prediction ability in 15°C T_night_ was comparable between the second set and the same number of randomly chosen SNPs from the third set (Supplementary Fig. 5A). This can be explained by the relatively modest -log₁₀ (*P*) values of most SNPs in the second set in 15°C Tₙᵢ_g_ₕₜ (≈ 4-5; Supplementary Table 3), which are not particularly higher than those of the third set (≥3.0). Inadequate correction for population structure can be ruled out to account for the high prediction ability with the third set, since no obvious clustering of accessions was observed in the PC analysis (Supplementary Fig. 6) and neither PC1 nor PC2 was correlated with the predicted F_q_’/F_m_’.

Hence, the high prediction ability achieved by the third set of SNPs most likely reflects their collective influence on midday F_q_’/F_m_’ in the accessions under the respective T_night_ conditions.

### SNPs associated with long-term response of F_q_’/F_m_’ to T_night_

While the third set of SNPs with -log_10_ (*P*) ≥ 3.0 gave equally high prediction abilities in both T_night_ conditions on all three days (Fig. 3C), the composition of associated SNPs was changing from day to day (Supplementary Fig. 7A). Nevertheless, part of the SNPs was repeatedly detected in 15°C T_night_, with larger overlaps found between two consecutive days (33‒58%) than between 16 and 18 DAS (29% and 24%). In total, 47 SNPs were associated with F_q_’/F_m_’ on all three days in 15°C T_night_. These 47 SNPs together explained nearly half of the F_q_’/F_m_’ variance in 15°C T_night_, each explaining 3 to 10% on different days (Table 1, Supplementary Table 6). The prediction based on the 47 SNPs was more accurate than using the first or the second set of SNPs, but not as good as the third set (Supplementary Table 7).

**Table 1:**
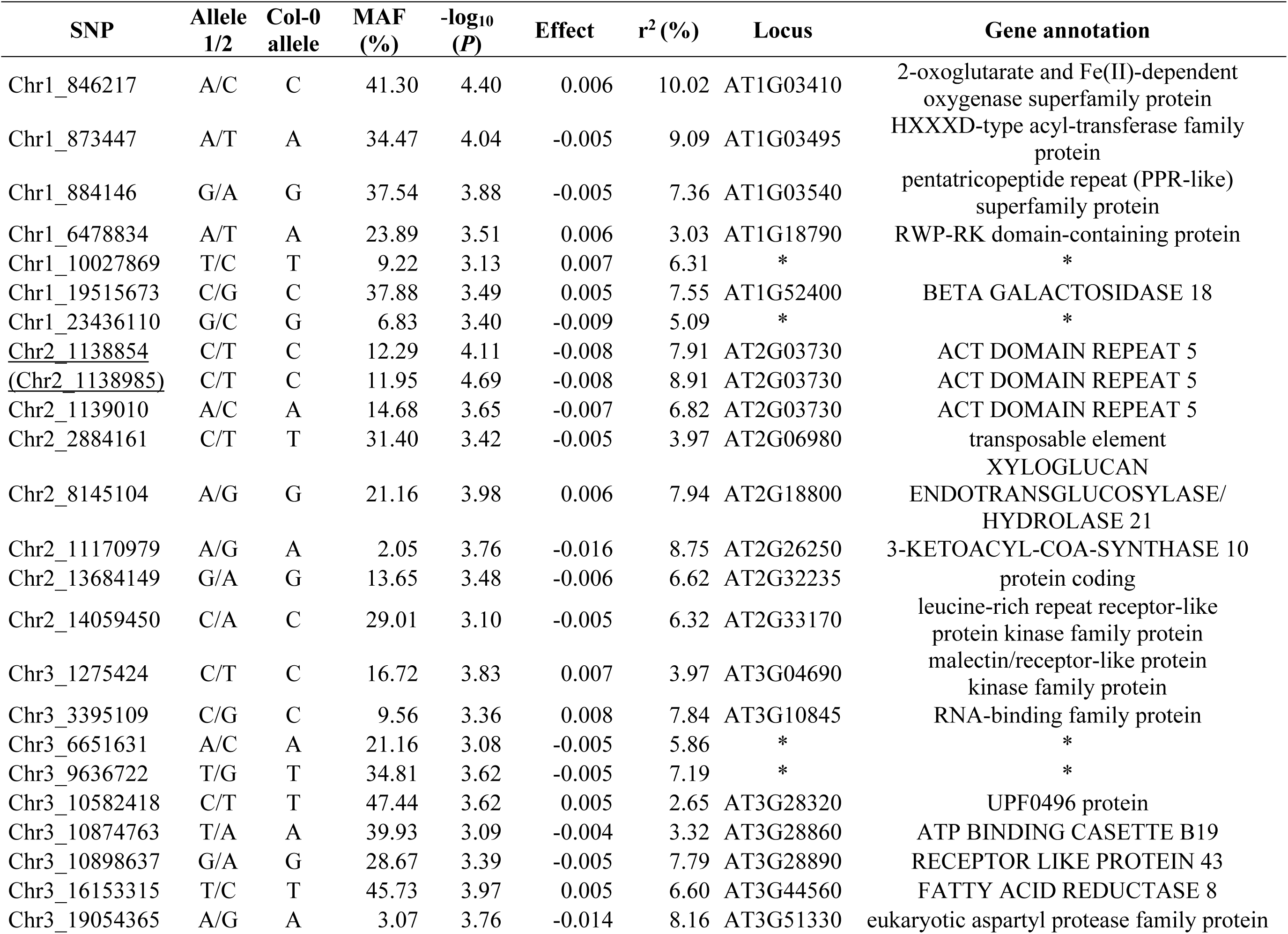
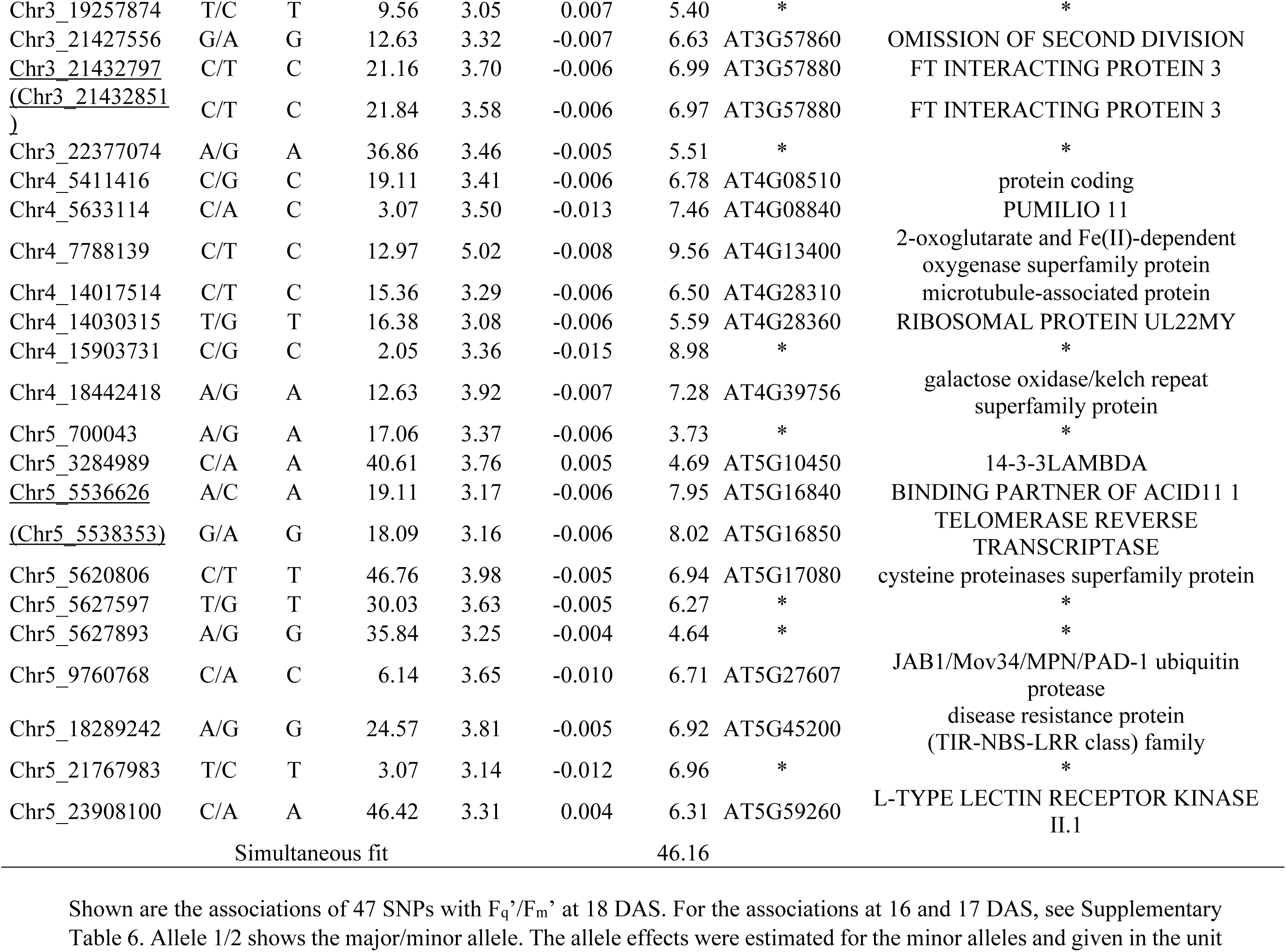
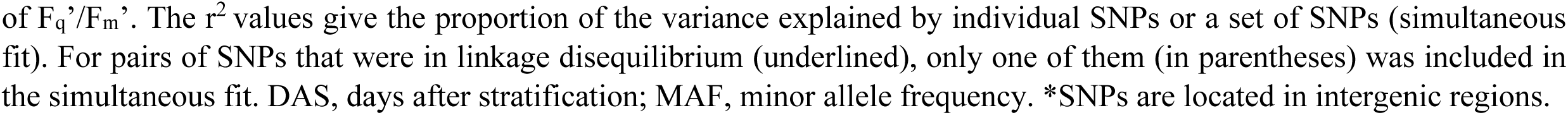
47 SNPs identified by GWAS with -log_10_ (*P*) ≥ 3.0 on three consecutive days in 15°C T_night_.

The 47 SNPs in 15°C T_night_ are likely to be identified by GWAS combining F_q_’/F_m_’ data of all three days. To verify this, adjusted entry-means of F_q_’/F_m_’ were computed for combined F_q_’/F_m_’ data, including measurement day as a random effect (Supplementary Fig. 8A). The heritability of these adjusted-entry means was 0.93, which is higher than the heritability calculated for each day separately (cf. Fig. 1E). The GWAS based on the combined data (Supplementary Fig. 8) confirmed the associations of 46 SNPs out of 47 (Supplementary Fig. 9). Further, 159 SNPs, which were associated with F_q_’/F_m_’ on one or two days in the separate analysis, were also detected by the combined analysis. In addition, 15 unique SNPs had - log_10_ (*P*) ≥ 3.0 in the combined analysis. A single SNP located in AT3G05050, encoding a protein kinase superfamily protein, passed the FDR threshold of 0.05 in the GWAS across three days (Supplementary Fig. 8B). This SNP was identified at 17 and 18 DAS in the separate analysis (SNP No. 7 in Fig. 2).

We consider the 47 SNPs in Table 1 as the core SNPs associated with F_q_’/F_m_’ variation among the Arabidopsis accessions in the 15°C T_night_ condition. None of these SNPs was found in loci known to be involved in photosynthesis. Besides the SNP in 3-KETOACYL-COA SYNTHASE 10 having the largest modulus effect of -0.016, five SNPs had the effect size between -0.01 and -0.015 (Table 1). Two of them are located in AT3G51330 and AT5G27607 which encode a eukaryotic aspartyl protease family protein and a JAB1/Mov34/MPN/PAD-1 ubiquitin protease, respectively. Another SNP was found in AT4G08840 coding for PUMILIO 11, a member of sequence-specific RNA-binding PUF proteins which regulate mRNA stability and translation^26^. The other two SNPs are located in intergenic regions, of which the one on chromosome 5 is within 1 kbp 5’-upstream of AT5G53580 encoding PYRIDOXAL REDUCTASE 1, an enzyme involved in the vitamin B6 salvage pathway^27^. The other one on chromosome 4 is within 300 bp 5’-upstream of AT4G32940 encoding a cysteine proteinase GAMMA VACUOLAR PROCESSING ENZYME that acts in the vacuole of vegetative organs^28^.

In striking contrast to 15°C T_night_, between-day overlaps of SNP-trait associations were no more than 2% to 6% in 20°C T_night_, as little as the overlaps between the two T_night_ conditions on each day (3-4%) (Supplementary Fig. 7). This pronounced day-to-day variation in 20°C T_night_ may signify dynamic nature of SNP-trait associations in this condition, with distinct sets of SNPs affecting midday F_q_’/F_m_’ on each day. Another possible explanation could be day-to-day fluctuation of -log_10_ (*P*) values around the chosen threshold, resulting in inclusion or exclusion of SNPs. If the former is the case, the SNPs identified at 16 or 17 DAS, for instance, will not allow accurate prediction of F_q_’/F_m_’ at DAS 18 (see the poor prediction ability of randomly chosen SNPs with -log_10_ (*P*) < 3.0 in Supplementary Fig. 4). In the latter case, on the other hand, we may find reasonably good predictions. Judging by the Pearson correlation coefficients between the actual and the predicted F_q_’/F_m_’, predictive performance for 18 DAS in 20°C T_night_ declined when the SNPs from 16 and 17 DAS in the same T_night_ condition or 18 DAS in 15°C T_night_ were used as predictors (Supplementary Table 8). Nonetheless, these correlation coefficients were substantially higher than using the first and the second sets of SNPs from the corresponding day and T_night_ (Supplementary Table 7). The SNPs from wrong days in the same 20°C T_night_ gave better predictions than the SNPs from the same day but in 15°C T_night_. Also, the SNPs from 17 DAS did better than those from 16 DAS for predicting F_q_’/F_m_’ at 18 DAS in 20°C T_night_ (Supplementary Table 8).

Taken together, these results imply that a greater number of loci than identified by -log_10_ (*P*) ≥ 3.0 on each day (Supplementary Table 4) were contributing to the F_q_’/F_m_’ variance among the Arabidopsis accessions, and that the associated loci were partly changing across days and T_night_ conditions. Between 15°C and 20°C T_night_, cooler nights led to a steady influence of a subset of associated SNPs on F_q_’/F_m_’, while no core set of SNPs was found in warmer nights.

### gBLUP outperforms ensemble methods in predicting F_q_’/F_m_’

The GPs described above were all realized by gBLUP. Given the large number of associated SNPs and their potential interactions, non-linear models may enable more accurate predictions than gBLUP. We therefore compared predictive performance of gBLUP with ensemble machine learning methods, namely, random forest^29,30^ and eXtreme Gradient Boosting (XGBoost)^31^. Both methods construct multiple individual models using different randomly sampled subsets of data. However, while random forest fits these models independently and in parallel, XGBoost fits them sequentially and iteratively, with each model correcting the errors of its predecessor^31^. We leveraged the optimized framework of XGBoost to handle the 211,771 SNPs for predicting F_q_’/F_m_’. Random forest was used for prediction by the GWAS-derived SNPs (i.e., the second and the third SNP sets). All models were trained by identical training sets and evaluated on the same validation sets to ensure comparability (Supplementary Fig. 10).

The ensemble methods performed similarly to gBLUP in predicting F_q_’/F_m_’ by the 211,771 SNPs and the second set of SNPs (Table 2). The prediction of XGBoost is unlikely to be affected by population structure because adjusting the genotypes for population structure did not improve the prediction (Supplementary Table 9). As seen by gBLUP, the information on the GWAS-derived SNPs improved predictive performance of the ensemble methods in both 15°C and 20°C T_night_ (Table 2). The best prediction was achieved by the third set of SNPs with -log_10_ (*P*) ≥ 3.0 in both T_night_ on all three days, whereby gBLUP outperformed random forest. Specifically, gBLUP produced higher correlations between the actual and predicted F_q_’/F_m_’, whereas random forest tended to overestimate in accessions with low F_q_’/F_m_’ and slightly underestimate in those with high F_q_’/F_m_’, yielding flatter regression slopes (Supplementary Fig. 11).

**Table 2:** Performance of gBLUP and ensemble methods in predicting F_q_’/F_m_’.

| $T_{\text{night}}$ | Method | DAS | Predictors | | | | | |
| --- | --- | --- | --- | --- | --- | --- | --- | --- |
| | | | 211,771 SNPs | | FDR threshold | | $-\log_{10}(P) \geq 3$ | |
|  |  |  | RMSE | COR | RMSE | COR | RMSE | COR |
| 15°C | gBLUP | 16 | 0.022±0.001 | 0.20±0.13 | 0.021±0.001 | 0.32±0.06 | 0.014±0.001 | 0.79±0.01 |
|  |  | 17 | 0.022±0.002 | 0.23±0.06 | 0.020±0.001 | 0.48±0.08 | 0.012±0.000 | 0.84±0.04 |
|  |  | 18 | 0.021±0.002 | 0.23±0.02 | 0.018±0.000 | 0.51±0.12 | 0.012±0.001 | 0.82±0.06 |
|  | random forest or XGBoost | 16 | 0.023±0.002 | 0.09±0.06 | 0.022±0.001 | 0.29±0.03 | 0.018±0.002 | 0.63±0.08 |
|  |  | 17 | 0.023±0.003 | 0.04±0.05 | 0.021±0.002 | 0.39±0.07 | 0.018±0.002 | 0.71±0.05 |
|  |  | 18 | 0.021±0.002 | 0.16±0.05 | 0.018±0.001 | 0.48±0.12 | 0.016±0.001 | 0.74±0.07 |
| 20°C | gBLUP | 16 | 0.022±0.001 | 0.15±0.09 | 0.016±0.001 | 0.69±0.07 | 0.010±0.001 | 0.89±0.02 |
|  |  | 17 | 0.018±0.001 | 0.25±0.01 | 0.015±0.001 | 0.62±0.01 | 0.009±0.000 | 0.86±0.03 |
|  |  | 18 | 0.016±0.001 | 0.21±0.03 | 0.013±0.001 | 0.59±0.08 | 0.008±0.001 | 0.86±0.03 |
|  | random forest or XGBoost | 16 | 0.022±0.001 | 0.22±0.10 | 0.018±0.001 | 0.60±0.07 | 0.017±0.001 | 0.71±0.02 |
|  |  | 17 | 0.017±0.002 | 0.27±0.06 | 0.015±0.001 | 0.54±0.06 | 0.013±0.001 | 0.72±0.04 |
|  |  | 18 | 0.016±0.001 | 0.30±0.04 | 0.014±0.001 | 0.51±0.09 | 0.013±0.001 | 0.65±0.02 |

### Validation experiment confirmed F_q_’/F_m_’ predictions in independent and genetically diverse accessions

Encouraged by the high accuracy of prediction using the GWAS-derived SNPs with -log_10_ (*P*) ≥ 3.0 (Fig. 3C; Table 2), we proceeded to examine whether the models, which were trained and tested on the 293 accessions in the GWAS panel, can predict F_q_’/F_m_’ for genetically diverse Arabidopsis accessions in 15°C and 20°C T_night_. Thus, the above-described gBLUP and random forest models with the third SNP set were applied to 1,014 accessions from the RegMap population^32^. The F_q_’/F_m_’ values predicted by gBLUP showed similar distributions in the 293 accessions and the 1,014 accessions under both T_night_ conditions (Supplementary Fig. 12). The predictions by the linear and non-linear methods were largely comparable for the 1,014 accessions, although random forest predicted higher F_q_’/F_m_’ values than gBLUP for accessions with low F_q_’/F_m_’ (Fig. 4A, B), as was seen during the cross-validation in the 293 accessions (Supplementary Fig. 11).

**Figure 4.**
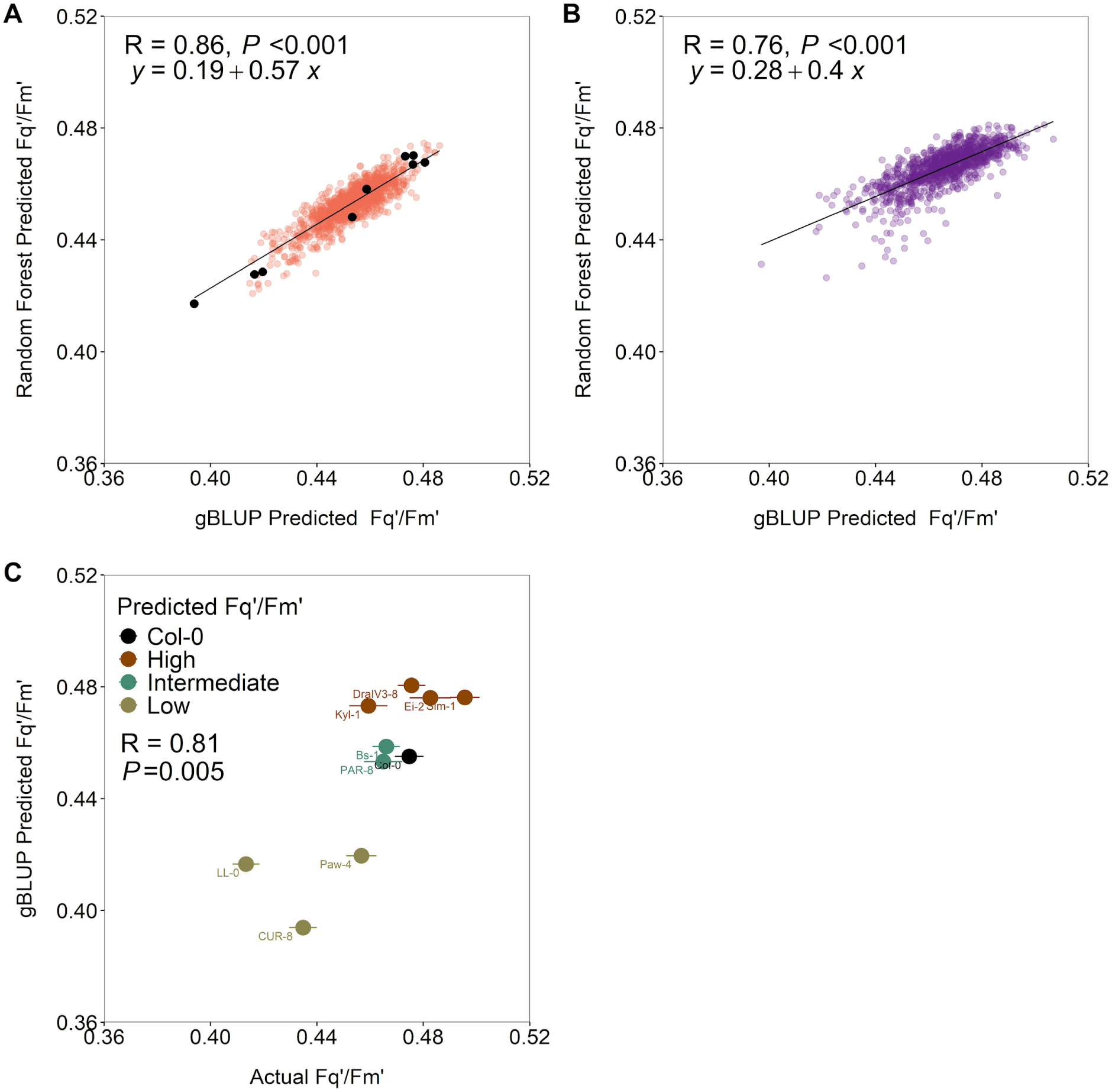
Genomic prediction of F_q_’/F_m_’ and experimental validation. Prediction in 1,014 non-phenotyped accessions using gBLUP or random forest based on the GWAS-derived SNPs with -log_10_ (*P*) ≥ 3.0 in 15°C (A) or 20°C (B) T_night_ at 18 days after stratification (DAS). Circles represent predicted F_q_’/F_m_’ of individual accessions. For nine accessions (marked with black circles in panel A; listed in Supplementary Table 10), prediction by gBLUP was experimentally validated in the 15°C T_night_ condition (C). Columbia-0 (Col-0) was included as reference in the validation experiment. Different colors of the accessions in panel C denote low, intermediate, and high predicted F_q_’/F_m_’ levels. Error bars are ±SE of the adjusted entry-means of actual F_q_’/F_m_’ of individual accessions. R indicates the Pearson correlation coefficient and *P* the significance. The number of replicate plants per accession was: Bs-1, n = 14; CUR-8, n = 19; DralIV3-8, n = 18; Ei-2, n = 3; Kyl-1, n = 4; LL-0, n = 25; PAR-8, n = 4; Paw-4, n = 11; Sim-1, n = 11; Col-0, n = 14. For validation at 16 and 17 DAS, see Supplementary Fig. 13.

We then conducted an experiment to validate the F_q_’/F_m_’ predictions. The 15°C T_night_ condition was chosen for the validation experiment since the correlation between gBLUP and random forest predictions was higher and the difference between the highest and the lowest predicted F_q_’/F_m_’ was larger in 15°C than in 20°C T_night_ (Fig. 4A, B), although heritability of F_q_’/F_m_’ was lower (Fig. 1E) and model predictions were less accurate in 15°C T_night_ (Table 2). Nine accessions having low, intermediate and high predicted F_q_’/F_m_’ (Fig. 4A; Supplementary Table 10) were grown alongside a reference genotype Col-0 and midday F_q_’/F_m_’ was measured at 16, 17 and 18 DAS. Overall, the predicted F_q_’/F_m_’ values were in very good agreement with the values observed in the accessions (R = 0.81 and *P* = 0.005 in Fig. 4C; Supplementary Fig. 13). While very small differences (ΔF_q_’/F_m_’ < 0.03) between high and intermediate accessions were difficult to verify experimentally, two accessions having the lowest F_q_’/F_m_’ (LL-0 and CUR-8) were correctly identified on all three days, supporting the models’ robustness and applicability across diverse populations.

## Discussion

Temperature profoundly influences biological processes. Up to an optimal point, rising temperatures accelerate biochemical and physiological reactions, whereas lower temperatures slow down enzyme activity and cellular processes, limiting growth and development. Reduced sink activity (i.e., the activity of a non-photosynthesizing organ or tissue to use or store photoassimilates) and the resulting sugar accumulation in leaves can trigger feedback suppression of photosynthesis^12,33^. In accordance, Arabidopsis plants grown in 15°C T_night_ exhibited smaller rosettes and lower midday F_q_’/F_m_’ than those in 20°C T_night_ (Fig. 1A, C). Since F_q_’/F_m_’ was measured under identical conditions for both T_night_ treatments, lower F_q_’/F_m_’ observed in 15°C T_night_ suggests feedback regulation of photosynthesis by sink limitation. Similar reductions in photosynthetic efficiency under low T_night_ have been reported in grapevine^8^, sugar beet^34^, and cotton^35^. Low T_night_ may also delay the development of photosynthetic capacity; the distribution of F_q_’/F_m_’ across accessions was comparable between DAS 18 in 15°C T_night_ and DAS 16 in 20°C T_night_ (Fig. 1A; Supplementary Table 2). Despite higher growth and midday F_q_’/F_m_’ in 20°C T_night_, predawn F_v_/F_m_ of the plants was slightly lower in 20°C than 15°C T_night_ (Fig. 1B), possibly reflecting a more reduced plastoquinone pool in the thylakoid membrane during warmer nights.

While both growth and PSII efficiency were influenced by T_night_ and genetic effects, F_q_’/F_m_’ stood out with the highest heritability of all parameters measured in this study (Fig. 1E). Broad-sense heritability represents the proportion of total phenotypic variance attributed to genetic variance, with higher values indicating stronger genetic control. In fact, the genetic variance component of F_q_’/F_m_’ was statistically significant (*P* < 0.001) on all three days in both 15°C and 20°C T_night_. Interestingly, heritability increased from DAS 16 to 18 for all parameters except RGR in 15°C T_night_ (Fig. 1E). Temporal changes in heritability ‒throughout the day, across developmental stages, and over growing seasons‒ have been reported for photosynthesis- and growth-related traits in Arabidopsis^36^, canola^37^, and barley^38^. The lower heritability observed at DAS16 is consistent with greater environmental sensitivity and higher ontogenetic variation in younger plants^39^, which often reduces heritability^40^. Another intriguing observation is the opposite effects of T_night_ on heritability of PSII efficiency and growth; F_q_’/F_m_’ and F_v_/F_m_ showed lower heritability in 15°C T_night_ whereas the heritability of projected rosette area and especially RGR was higher in this condition (Fig. 1E). Lower heritability of PSII efficiency in 15°C T_night_ may be due to slower development and/or higher environmental sensitivity of photosynthesis in this condition. Higher heritability of growth in 15°C T_night_, on the other hand, is difficult to explain by development. Cooler nights may creat a more selective environment for growth, amplifying genetic differences among accessions that are less pronounced in warmer T_night_.

Our GWAS revealed another distinct effect of cooler nights on the genetic control of F_q_’/F_m_’, manifested as the steady contributions of a subset of associated SNPs (Table 1, Supplementary Table 6). None of the 47 core SNPs identified in 15°C T_night_ is located in loci that are known to be involved in photosynthesis. This finding is in line with the previous GWAS study in Arabidopsis, reporting the absence of photosynthetic genes among 63 candidates that were associated with high-light response of F_q_’/F_m_’ (Φ_PSII_ in that study)^41^. Contrary to the picture in 15°C T_night_, only a few SNPs were repeatedly identified on different days in 20°C T_night_ (Supplementary Fig. 7). Since gBLUP showed fairly good predictive performance using SNPs from wrong days for this T_night_ condition (Supplementary Table 8), much better than using all 211,771 SNPs (Table 2; Supplementary Table 7) and randomly selected less significant SNPs (Supplementary Fig. 4), it is unlikely that 20°C T_night_ caused radical day-to-day shifts in the genetic architecture of F_q_’/F_m_’. Rather, it seems to reflect the existence of numerous small-effect loci, far beyond the 160-260 SNPs identified by -log_10_ (*P*) ≥ 3.0 on each day (Supplementary Table 4). The higher predictive performance obtained by the SNPs from the same T_night_ than the other T_night_, and from the previous day than two days earlier (Supplementary Table 8), underscores the roles of T_night_ and development in shaping the genetic basis of F_q_’/F_m_’.

The multitude of small-effect variants associated with F_q_’/F_m_’ aligns with the omnigenic model^42^ which posits that any gene expressed in relevant cells can indirectly influence a complex trait by modulating the expression or activity of primary genes having direct effects. Photosynthesis may exemplify this principle; it relies on many enzymes and cofactors whose synthesis, regulation, and degradation depend on an extensive network of other enzymes, transcription factors, and transporters, themselves subject to similar control. Close interconnections between photosynthesis, central metabolism, and growth further strengthen the omnigenic-like nature of photosynthesis. The very fact that midday F_q_’/F_m_’ and its genetic architecture changed in response to growth T_night_, partly via sink regulation, highlights the inherent complexity of this trait. Distributing the control over many enzymes allows metabolic flux, like in photosynthesis and related pathways, to be regulated efficiently with minimal disruption and perturbation^43^. This system-level control arises from synergistic interactions among multiple control sites‒ a concept that may also apply to the genetic control of complex traits where numerous small-effect loci collectively influence phenotypic outcomes under variable environmental conditions and at different developmental stages.

When a trait is highly polygenic, how can SNP-trait associations be meaningfully evaluated? Assessing a limited number of loci by mutant characterization seems insufficient, since individual SNPs exert only marginal effects and collective influence of the majority is neglected. We therefore adopted a holistic approach that focuses on the combined contribution of many SNPs, considering SNP-trait associations to be reliable if they yield significant improvements in the performance of GP models. The results demonstrated that the GWAS-derived SNPs, particularly those selected by -log_10_ (*P*) ≥ 3.0, substantially and consistently enhanced the prediction of F_q_’/F_m_’ in the 293 accessions (Fig. 3C) compared to using all (Fig. 3A; Table 2; Supplementary Table 7) or random SNPs (Suppplementary Fig. 4). The prediction was improved regardless of whether linear or non-linear models were applied, although gBLUP outperformed random forest (Table 2; Supplementary Fig. 11). Increases in GP accuracy following the incorporation of GWAS results have been reported in different plant species such as rice^44^, maize^45^, wheat^46^, and poplar^47^. Without GWAS-derived information (i.e., using all 211,771 SNPs), the median prediction ability of gBLUP for F_q_’/F_m_’ was ∼0.33 (Fig. 3A), similar to the previous study in which gBLUP achieved the accuracy of ∼0.3 and ∼0.4 for predicting Φ_PSII_ (F_q_’/F_m_’) in 344 Arabidopsis accessions under light intensities of 100 and 550 μmol photons m^-2^ s^-1^, respectively^48^. Neither Bayes A nor Bayes B improved the predictive performance (Supplementary Table 5), despite their capacity to model heterogenous marker effects^16^. This agrees with previous findings that gBLUP and Bayes methods performed similarly when traits were governed by numerous small-effect loci^49,50^. The limited accuracy achieved by genome-wide SNPs could be due to the introduction of noise by the vast majority of non-associated SNPs. Conversely, small sets of SNPs identified by stringent thresholds (-log_10_ (*P*) ≥ FDR and/or -log_10_ (*P*) > 4.0 on at least two days) yielded rather modest and inconsistent gains in prediction ability (Fig. 3B; Table 2; Supplementary Table 7), suggesting that these SNPs alone cannot sufficiently account for the observed variation in F_q_’/F_m_’ across the accessions. Hence, conventional reliance on conservative thresholds may not be good practice for GWAS on complex traits underpinned by numerous small-effect loci when aiming to maximize the prediction ability. A more effective strategy could be to systematically assess multiple significance thresholds and identify the sets of associated variants that allow high prediction accuracy.

Whilst predictions can be significantly improved by incorporating GWAS-derived information, the chief purpose of GP is to predict the potential of individuals based on their genome-wide marker data alone. Yet, GP models may not transfer well across genetically diverse populations, as prediction ability is strongly influenced by kinship, population structure, and the extent of linkage disequilibrium^51^. Thus, to evaluate cross-population applicability, we tested the GWAS-based models, which were trained on the 293 phenotyped accessions from the HapMap population^22^, against 1,014 non-phenotyped accessions from the RegMap population^32^. Using the SNPs with -log_10_ (*P*) ≥ 3.0 as predictors, the predictions of gBLUP and random forest were strongly and linearly correlated in the 1,014 accessions under both 15°C and 20°C T_night_ (Fig. 4A, B). To further assess model performance, we experimentally validated the predictions in a subset of accessions spanning a range of predicted F_q_’/F_m_’ values (Fig. 4A; Supplementary Table 10). Albeit rarely performed in GP studies, experimental validation offers a definitive assessment of model performance in a given context. Our results confirmed the GP models’ capacity to correctly identify genotypes with low F_q_’/F_m_’ (LL-0 and CUR-8) in the 15°C T_night_ condition, whereas minimal differences between high and intermediate genotypes could not be verified by the measurements (Fig. 4C; Supplementary Fig. 13). Accordingly, the GWAS-derived SNPs seem to capture sufficient genetic information to reliably identify low-F_q_’/F_m_’ individuals in an independent, genetically diverse Arabidopsis population under our growth conditions. Moreover, we noticed that a previous study^52^, which had been conducted under conditions distinct from ours, also reported low Φ_PSII_ (F_q_’/F_m_’) in LL-0, suggesting a certain degree of transferability of the models even across environments.

In summary, our findings uncover the highly polygenic architecture of photosynthesis and its differential responses to warmer *vs* cooler nights. The results provide a framework for leveraging GWAS and GP to explore complex quantitative traits, such as photosynthesis, toward breeding crops that are resilient to future climate challenges.

## Materials and Methods

### Plant materials and growth conditions

The accessions of *Arabidopsis thaliana* (L.) Heynh. (Supplementary Table 1) from the global HapMap population^22^ were purchased from the Nottingham Arabidopsis Stock Centre (N76309). Seeds were sown automatically in 576-cell trays containing moist substrate (type Topf 1.5; Balster Einheitserdewerk, Fröndenberg, Germany) with the automated PhenoSeeder system^53^. After three to four days of stratification at 4°C in the dark, the trays with seeds were transferred to a climate chamber running with 12 h/12 h light/dark cycle at 26°C T_day_, 20°C or 15°C T_night_, and constant 60% relative air humidity. The illumination was provided by white LEDs (XLamp CXA2520; Cree LED, Durham, NC, USA). The diurnal light intensity regime was as follows: 4 h of linear increase to the maximum intensity of ∼600 µmol photons m^-2^ s^-1^ (measured at the plant height), then 4 h of constant light at this intensity, followed by 4 h of linear decrease. The daily light integral was ∼18.7 mol photons m^-2^ d^-1^.

Germination was monitored daily at around midday (4.5 - 7.5 h into the light period) for eight days after stratification using an automated image-based device^54^. At 11 DAS, seedlings were transferred to pots (7 x 7 x 7 cm; one plant per pot) filled with substrate (Lignostrat Dachgarten extensiv; HAWITA, Vechta, Germany) and placed under the aforementioned conditions in the climate chamber. Pots were randomly placed in plastic inlets (24 pots per inlet; Nr.68124; Bachmann Plantec, Hochdorf, Switzerland) where they stayed throughout the experiments (Supplementary Fig. 14). Accession*treatment*replication combinations were evaluated in six independent experiments (three experiments per treatment). The accession Col-0 was included in each experiment as reference, with six and ten replicate plants in total at 15°C and 20°C T_night_, respectively. Plants were well-watered throughout cultivation and experiments.

An additional experiment was conducted using nine selected accessions (Bs-1, CUR-8, DralIV3-8, Ei-2, Kyl-1, LL-0, PAR-8, Paw-4 and Sim-1; Supplementary Table 10) and Col-0 (reference) to validate the model prediction of F_q_’/F_m_’ in an independent set of accessions. Seeds of these accessions were purchased from the Nottingham Arabidopsis Stock Centre and treated in the same way as described above. Plants were placed in the plastic inlets under the 15°C T_night_ conditions. The number of replicate plants used in the validation experiment was: Bs-1, n = 14; Col-0, n = 15; CUR-8, n = 19; DraIV3-8, n = 18; Ei-2, n = 3; Kyl-1, n = 4; LL-0, n = 27; PAR-8, n = 4; Paw-4, n = 13; Sim-1, n = 11.

### Rosette growth analysis

Projected rosette area was measured at 16, 17, 18 and 19 DAS using the image-based GROWSCREEN method^55^. The measurements were performed at around midday (4.5 - 8 h into the light period) when leaves were positioned almost horizontally. Relative growth rate (RGR, % d^-1^) was calculated for each plant using the projected rosette area measured on two consecutive days, as previously described^56^.

### Chlorophyll *a* fluorescence analysis

Chlorophyll *a* fluorescence measurements were performed in dark-adapted and light-adapted plants under the growth conditions in the climate chamber at 16, 17 and 18 DAS. Dark-adapted plants were measured during the night (8 - 11.5 h into the dark period) at the respective T_night_ (20°C or 15°C). Light-adapted plants were measured around midday (4.5 - 8 h into the light period) when the light intensity stayed at ca. 600 μmol photons m^-2^ s^-1^ and the air temperature was 26°C. Measurements were performed using a LIFT instrument (model LIFT-REM 1.0; Soliense Inc., Shoreham, New York, USA) mounted on a tripod (Advanced VX mount; Celestron, Torrance, USA) with an automated positioning system. The plastic inlets with 24 plants were placed one by one in the designated position in front of the LIFT instrument (Supplementary Fig. 14). The distance between the LIFT instrument and the plants was between 63.5 and 71.8 cm depending on the position of the plants inside the inlet. For each plant, this distance remained unchanged throughout the experiment.

We used the LIFT excitation protocol FRRF_0.75ms_^57^ which is based on the fast repetition rate approach^58^. This protocol employs an excitation flash consisting of 300 sub-saturating short (1,6 μs) pulses (“flashlets”) triggered at a 2.5-μs interval for 0.75 ms to reduce the primary quinone acceptor of PSII (Q_A_), followed by 125 flashlets triggered at exponentially increasing intervals for 209 ms to monitor Q_A_ reoxidation. The excitation flashes were applied by four LED channels (445, 470, 505, and 535 nm) with the average intensity in the Q_A_ reduction phase of 21,890 and 18,050 µmol photons m^−2^ s^−1^ (determined according to ^59^ at a distance of 60 cm from the LIFT instrument), respectively, in the GWAS experiments and the validation experiment. Emission of chlorophyll *a* fluorescence was detected at 685 (± 10) nm. Measurements in dark-adapted plants were performed with a single flash, while in light-adapted plants five flashes were applied with a 1.5-s interval to obtain a mean value. The light-adapted measurements were started at least 30 s after placing the inlet in front of the LIFT.

The minimal (F_o_), the maximal (F_m_ and F_m_’) and steady-state (F_s_) fluorescence yield measured by the LIFT protocol^59^ were used to calculate the maximal (F_v_/F_m_ = (F_m_ – F_o_)/F_m_) and effective quantum yield of PSII (F_q_’/F_m_’ = (F_m_’ – F_s_)/F_m_’) in the dark-adapted and light-adapted states, respectively.

### Estimation of adjusted entry-means

Chlorophyll fluorescence and growth data were discarded when i) plants exhibited severely impaired growth, ii) plants were damaged by handling during the experiments, iii) accessions had rosette architecture that interfered with 2D image analysis, iv) accessions had only one replicate plant, or v) plants were classified as outliers in the statistical analyses. Data from 308 accessions remained after these cleaning steps. Linear mixed models were fitted to account for spatial heterogeneity of the growth and LIFT measurement conditions (Supplementary Fig. 14).

The models used ‒ with respective transformation of the response if required ‒ are:

Model 1: Adjust entry-means of projected rosette area

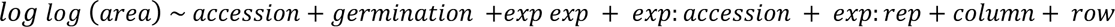

Model 2: Adjust entry-means of rosette RGR

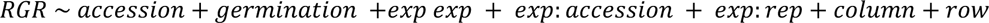

Model 3: Adjust entry-means of F_v_/F_m_

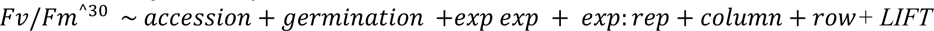

Model 4: Adjust entry-means of F_q_’/F_m_’

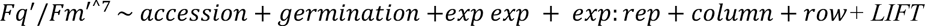

where *germination* indicates the day on which seedlings germinated, *exp* the experiment number (#1 to #3 for each T_night_ treatment), *rep* the plant replicate in each experiment, *column* and *row* the x and y position within the inlet (accounting for variations in growth conditions; Supplementary Fig. 14), and *LIFT* the plant position during the LIFT measurements (accounting for variations in the distance and angle between the LIFT instrument and the plants during the measurements; Supplementary Fig. 14). The effect of accessions was fixed, while germination date, experiment, accession nested in experiment, replicate nested in experiment, x-y positions, and LIFT measurement positions were random. Adjusted entry-means of F_q_’/F_m_’ across three days combined in the 15°C T_night_ condition were obtained by using Model 4 with day as an additional random effect.

The model used for the validation experiment with ten accessions is:

Model 5: Adjust entry-means of F_q_’/F_m_’

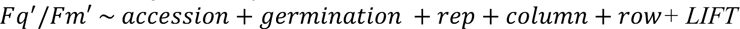

The models were fitted separately for each T_night_ treatment and measurement day using the *lmer* function in the *lme4* package of R^60^ (version 4.2.2). Adjusted entry-means were obtained using the function *emmeans* in the *emmeans* package. The model assumptions (i.e., normal distribution of residuals and homogeneity of variance) were verified for each model. Statistically significant differences between the T_night_ treatments were tested by two-tailed Student t-test (equal or unequal variance checked by F-test) using the *t.test* function. Broad-sense heritability, defined as the proportion of genetic variance to phenotypic variance, was estimated separately for each T_night_ condition and measurement day according to ^61^ using Models 1-4 with accession as random effect.

### Genome-wide association

Of the 308 accessions in the phenotyping experiments, 15 (627ME-4Y1, BUI, CLE-6, DraIV1-7, Gul1-2, Kas-2, NC-6, PHW-34, PUZ24, RRS, Sav-1, TOU-A1-12, TOU-A1-67, UKCW06202 and Wil-1) lacked SNP data and, thus, were excluded from GWAS. The SNP information was obtained from the regional mapping panel (RegMap^29^) at https://easygwas.biochem.mpg.de (method 75, accessed on 23.03.2023). The SNP dataset contained 214,051 variants, which were reduced to 211,771 variants after filtering out minor allele frequency (< 2%) using PLINK^62^ version v1.90b6.21. The ID numbers of the accessions (listed in Supplementary Table 1) were obtained from https://bergelsonlab.org/resources/a-thaliana/ (accessed on 22.03.2023). When the ID numbers were inconsistent between the SNP dataset and the accession list (Bg-2, Br-0, Col-0, Got-7 and Oy-0), information was obtained from http://naturalsystems.uchicago.edu/naturalvariation/hapmap/ (accessed on 23.03.2023).

The analysis of SNP-trait association was performed separately for each T_night_ condition and measurement day by GAPIT (version 3) using FarmCPU model^63^ based on the adjusted entry-means of F_q_’/F_m_’. Population structure was accounted for by using up to five PCs (Supplementary Fig. 2). To select the appropriate number of PCs, we performed PC analysis based on the 211,771 SNPs (Supplementary Fig. 1) and fitted the linear models to F_q_’/F_m_’ by adding an increasing number of PCs as covariates with the *sommer* package. The PCs that most explained the variance in F_q_’/F_m_’, assessed as the correlation coefficient based on log likelihood^64^, were included in the FarmCPU. Gene annotation was retrieved from TAIR10 (https://www.arabidopsis.org/).

The proportion of phenotypic variance explained by individual SNPs or sets of SNPs was estimated by fitting the linear models with SNPs as fixed effects using the *lm* function in the *stats* package of R. When pairs of SNPs were in linkage disequilibrium (R^2^ > 0.8), only one of them was included in a set of SNPs (simultaneous fit). The proportion of explained phenotypic variance was assessed as the squared correlation coefficient (r^2^) between the actual and predicted F_q_’/F_m_’ in five-fold cross-validation using 40 replications. Effects of the SNP-trait associations detected in the full dataset were estimated newly in each run of the cross-validation. For this purpose, phenotypic data were randomly split into five groups, of which four comprised the training set and the remaining group served as the test set. The models were then trained on the training set and evaluated on the test set. This procedure was repeated five times using each of the five groups as the test set.

### Genomic prediction

Genomic prediction of F_q_’/F_m_’ was performed using gBLUP^18^ and two ensemble machine learning methods, namely random forest^29,30^ and XGBoost^31^. We additionally tested Bayesian models BayesA and BayesB^16^ to verify whether prediction ability based on the 211,771 SNPs can be improved by assuming non-homogenous effects of SNPs. Prediction models were fitted to the adjusted entry-means of F_q_’/F_m_’ separately for each T_night_ treatment and measurement day. Three different SNP sets were used as predictors: i) all 211,771 SNPs used in the GWAS, ii) the GWAS-derived SNPs with -log_10_ (*P*) ≥ FDR threshold (0.05) and/or -log_10_ (*P*) > 4.0 on at least two measurement days, and iii) the GWAS-derived SNPs with -log_10_ (*P*) ≥ 3.0 on each day.

Genomic prediction based on gBLUP was performed with the *sommer* package of R. Prediction ability was estimated as the median Pearson correlation coefficient between the actual and the predicted F_q_’/F_m_’ across all five-fold cross-validation runs. BayesA and BayesB were used as implemented in the *BGLR* package^65^ based on the 211,771 SNPs. Gibbs sampling was performed with 30,000 iterations, of which 10,000 samples were discarded as burn-in. Prediction using random forest was based on the GWAS-derived SNPs, namely, the ones with -log_10_ (*P*) ≥ FDR (0.05) and/or -log_10_ (*P*) > 4.0 on at least two measurement days, and those with -log_10_ (*P*) ≥ 3.0 on each day. The prediction was performed by the *caret* package^66^ of R using 500 trees and 5-fold cross-validation. The number of features selected as predictors in each tree node was *p*-1, where *p* is the total number of SNPs in the set.

Genomic prediction by XGBoost based on the 211,771 SNPs was performed on the JUWELS-Booster high-performance computing system (https://apps.fz-juelich.de/jsc/hps/juwels/booster-overview.html) of Forschungszentrum Jülich^67^. We used XGBoost as part of the Scikit-Learn library^68^ in Python 3.10.4 (https://www.python.org). To identify the combination of hyperparameters that maximized the predictive performance of XGBoost, we performed a grid search (Supplementary Table 11) with mean squared error as the loss function. Since the grid search over maximal depth of tree and number of estimators yielded comparable predictive performance, the prediction results are presented for the grid search over maximal depth of tree (3 or 5) and 100 trees. Correction for population structure was performed by taking the residuals of 250 PCs from a PC analysis of the genotypes^69^ as the inputs for prediction by XGBoost. Since similar prediction of F_q_’/F_m_’ was obtained with 100 or 500 trees and with 3, 5, 6 or 10 maximum tree depth as described above, grid search was performed over the maximum depth (3 or 5) with 100 estimators.

The predictive performance of gBLUP and random forest (based on the GWAS-derived SNPs) or gBLUP and XGBoost (211,771 SNPs) was compared by using the same data sets for training as well as for testing. Phenotypic data of the 293 accessions were randomly split into training and test sets at 4:1 ratio and three independent replicates were run. Predictive performance was assessed as the prediction ability (as described above) and root mean square error (RMSE) of the test sets. The F_q_’/F_m_’ data of all 293 accessions were used for training and testing the models at 17 and 18 DAS, whereas one accession (T1080) was excluded at 16 DAS.

The prediction models trained and tested on F_q_’/F_m_’ of the 293 accessions were used to predict F_q_’/F_m_’ for 1.014 genetically diverse, non-phenotyped accessions from the RegMap population^29^ using gBLUP and random forest based on the GWAS-derived SNPs with -log_10_ (*P*) ≥ 3.0. Nine accessions (Supplementary Table 10) having low, intermediate, and high predicted F_q_’/F_m_’ values (Fig. 4A) were selected and used for experimental validation of the model prediction in 15°C T_night_. Col-0 was included as reference in the validation experiment.

## Acknowledgements

The work of A.C.S.S was funded by the Deutsche Forschungsgemeinschaft (DFG), project ID 391465903/GRK 2466. S.M. and B.S. were supported by DFG under Germany’s Excellence Strategy – EXC-2048/2 – project ID 390686111. The GP with machine learning was supported by the Helmholtz Association Initiative and Networking Fund in the frame of Helmholtz AI. We thank the Jülich Supercomputing Centre for the access to JUWELS. A.C.S.S. and S.M. are grateful for valuable discussions with, and friendly support by Zbigniew Kolber (Soliense Inc.) during LIFT experiments. A.C.S.S. also thank Yuxi Niu, Laura Jansen and Elisa Sophie Großmann (IBG-2, FZJ) for their help during automated seed sowing.

## Author contributions

S.M. conceived the study. A.C.S.S., B.S., and S.M. designed the experiments. A.C.S.S conducted the experiments. A.C.S.S. analyzed the data aided by B.S. for GWAS and gBLUP, and by A.B. and S.K. for ensemble methods. A.F. provided technical support for image-based growth analysis and automated seed sowing. A.C.S.S. and S.M. wrote the manuscript with contributions of all coauthors.

