## Supplementary Information for "Highly polygenic control of photosynthetic responses to nighttime temperature in Arabidopsis studied by genomic prediction"

**Supplementary Table 1.** List of 308 *Arabidopsis thaliana* accessions.

Ecotype ID refers to the accession identification in the genotype data. Country refers to the origin of the accession.

| Accession | Ecotype ID | Country |
|---|---|---|
| 11ME1.32 | 8610 | United States of America |
| 11PNA4.101 | 8796 | United States of America |
| 328PNA054 | 8692 | United States of America |
| 627ME-4Y1 | — | United States of America |
| Aa-0 | 7000 | Germany |
| Ag-0 | 6897 | France |
| Alc-0 | 6988 | Spain |
| ALL1-2 | 1 | France |
| Amel-1 | 6990 | Netherlands |
| An-1 | 6898 | Belgium |
| An-2 | 6996 | Belgium |
| Ann-1 | 6994 | France |
| App1-16 | 5832 | Sweden |
| Ba-1 | 7014 | United Kingdom |
| Ba1-2 | 8256 | Sweden |
| Baa-1 | 7002 | Netherlands |
| Bay-0 | 6899 | Germany |
| Be-1 | 7011 | Germany |
| Belmonte-4-94 | 957 | Italy |
| Benk-1 | 7008 | Netherlands |
| Bg-2 | 6709 | United States of America |
| Bla-1 | 8264 | Spain |
| Blh-1 | 8265 | Czech Republic |
| Blh-2 | 7035 | Czech Republic |
| Boot-1 | 7026 | United Kingdom |
| Bor-4 | 6903 | Czech Republic |
| Br-0 | 6904 | Czech Republic |
| Bro1-6 | 8231 | Sweden |
| Bs-2 | 7004 | Switzerland |
| Bsch-0 | 7031 | Germany |
| BUI | — | France |
| Bu-0 | 8271 | Germany |
| Bur-0 | 6905 | Ireland |
| C24 | 6906 | Portugal |
| Ca-0 | 7062 | Germany |
| CAM-16 | 23 | France |
| CAM-61 | 66 | France |
| Can-0 | 8274 | Spain |
| Cen-0 | 8275 | France |
| Cha-0 | 7069 | Switzerland |

| | | |
|---|---|---|
| Chat-1 | 7071 | France |
| CIBC-17 | 6907 | United Kingdom |
| CIBC2 | 6727 | United Kingdom |
| CIBC4 | 6729 | United Kingdom |
| CIBC5 | 6730 | United Kingdom |
| Cit-0 | 7075 | France |
| CLE-6 | — | France |
| Co-2 | 7078 | Portugal |
| Co-4 | 7080 | Portugal |
| Col-0 | 6909 | United States of America |
| CSHL-5 | 6744 | United States of America |
| Ct-1 | 6910 | Italy |
| CUR-3 | 81 | France |
| Cvi-0 | 6911 | Cape Verde |
| Da-0 | 7094 | Germany |
| Db-0 | 7100 | Germany |
| Di-1 | 7098 | France |
| Do-0 | 7102 | Germany |
| DraIV1-14 | 5896 | Czech Republic |
| DraIV1-5 | 5887 | Czech Republic |
| DraIV1-7 | — | Czech Republic |
| DraIV6-16 | 5987 | Czech Republic |
| DraIV6-35 | 6005 | Czech Republic |
| Duk | 6008 | Czech Republic |
| Ede-1 | 7110 | Netherlands |
| Eden-2 | 6913 | Sweden |
| Edi-0 | 6914 | United Kingdom |
| Ep-0 | 7123 | Germany |
| Es-0 | 7126 | Finland |
| Est-0 | 7128 | Russia |
| Est-1 | 6916 | Russia |
| Fab-4 | 6918 | Sweden |
| Fei-0 | 8215 | Portugal |
| Fi-1 | 7139 | Germany |
| Fja1-2 | 6019 | Sweden |
| Fja1-5 | 6020 | Sweden |
| Fr-4 | 7135 | Germany |
| Ga-0 | 6919 | Germany |
| Ga-2 | 7141 | Germany |
| Gd-1 | 8296 | Germany |
| Ge-0 | 8297 | Switzerland |
| Ge-1 | 7145 | Switzerland |
| Gel-1 | 7143 | Netherlands |
| Gie-0 | 7147 | Germany |

| | | |
|---|---|---|
| Gö-0 | 7151 | Germany |
| Got-7 | 6921 | Germany |
| Gr-1 | 8300 | Austria |
| Gul1-2 | — | Sweden |
| Gy-0 | 8214 | France |
| Ha-0 | 7163 | Germany |
| Hau-0 | 7164 | Denmark |
| Hey-1 | 7166 | Netherlands |
| Hi-0 | 8304 | Netherlands |
| Hn-0 | 7165 | Germany |
| Hod | 8235 | Czech Republic |
| Hov4-1 | 8306 | Sweden |
| Hovdala-2 | 6039 | Sweden |
| HR-5 | 6924 | United Kingdom |
| Hs-0 | 8310 | Germany |
| HSm | 8236 | Czech Republic |
| In-0 | 8311 | Austria |
| JEA | 91 | France |
| Jl-3 | 7424 | Czech Republic |
| Ka-0 | 8314 | Austria |
| Kas-2 | — | India |
| KBS-Mac-8 | 1716 | United States of America |
| Kelsterbach-2 | 7188 | Germany |
| Kelsterbach-4 | 8420 | Germany |
| Kin-0 | 6926 | United States of America |
| Kl-5 | 7199 | Germany |
| Kn-0 | 7186 | Lithuania |
| Kno-18 | 6928 | United States of America |
| Koln | 8239 | Germany |
| Kr-0 | 7201 | Germany |
| Kro-0 | 7206 | Germany |
| Krot-2 | 7205 | Germany |
| Kulturen-1 | 8240 | Sweden |
| LAC-3 | 94 | France |
| LAC-5 | 96 | France |
| LDV-14 | 104 | France |
| LDV-25 | 116 | France |
| LDV-58 | 149 | France |
| Li-3 | 7224 | Germany |
| Li-5:2 | 7227 | Germany |
| Li-7 | 7231 | Germany |
| Liarum | 8241 | Sweden |
| Lillo-1 | 8242 | Sweden |
| LI-OF-095 | 641 | United States of America |

| | | |
|---|---|---|
| Lip-0 | 8325 | Poland |
| Lis-1 | 8326 | Sweden |
| Lis-2 | 8222 | Sweden |
| Lisse | 8430 | Netherlands |
| Lm-2 | 8329 | France |
| Lom1-1 | 6042 | Sweden |
| Lov-5 | 6046 | Sweden |
| Lp2-2 | 7520 | Czech Republic |
| Lp2-6 | 7521 | Czech Republic |
| Lund | 8335 | Sweden |
| Lz-0 | 6936 | France |
| Map-42 | 2057 | United States of America |
| Mc-0 | 7252 | United Kingdom |
| Mh-0 | 7255 | Poland |
| MIB-15 | 166 | France |
| MIB-22 | 173 | France |
| MIB-28 | 178 | France |
| MIB-84 | 223 | France |
| MNF-Che-2 | 1925 | United States of America |
| MNF-Jac-32 | 1967 | United States of America |
| MNF-Pot-48 | 1859 | United States of America |
| MNF-Pot-68 | 1867 | United States of America |
| Mnz-0 | 7244 | Germany |
| MOG-37 | 242 | France |
| Mr-0 | 7522 | Italy |
| Mrk-0 | 6937 | Germany |
| Mt-0 | 6939 | Libya |
| Mz-0 | 6940 | Germany |
| N13 | 7438 | Russia |
| N4 | 7446 | Russia |
| Na-1 | 8343 | France |
| Nc-1 | 7430 | France |
| NC-6 | — | United States of America |
| Nd-1 | 6942 | Switzerland |
| NFA-10 | 6943 | United Kingdom |
| NFA-8 | 6944 | United Kingdom |
| No-0 | 7275 | Germany |
| Nok-1 | 7270 | Netherlands |
| Nw-0 | 7258 | Germany |
| Nw-2 | 7260 | Germany |
| Nz1 | 7263 | New Zealand |
| Old-1 | 7280 | Germany |
| Omo2-1 | 7518 | Sweden |
| Or-0 | 7282 | Germany |

| | | |
|---|---|---|
| Or-1 | 6074 | Sweden |
| Ors-1 | 7283 | Romania |
| Ors-2 | 7284 | Romania |
| Ost-0 | 8351 | Sweden |
| Oy-0 | 6946 | Norway |
| Pa-2 | 7291 | Italy |
| PAR-3 | 258 | France |
| PAR-4 | 259 | France |
| PAR-5 | 260 | France |
| Paw-3 | 2150 | United States of America |
| Pent-1 | 2187 | United States of America |
| Per-1 | 8354 | Russia |
| Petergof | 7296 | Russia |
| PHW-10 | 7479 | United Kingdom |
| PHW-13 | 7482 | United Kingdom |
| PHW-14 | 7483 | United Kingdom |
| PHW-20 | 7490 | United Kingdom |
| PHW-26 | 7496 | United Kingdom |
| PHW-28 | 7498 | United Kingdom |
| PHW-33 | 7504 | Netherlands |
| PHW-34 | — | France |
| PHW-35 | 7506 | France |
| PHW-37 | 7508 | France |
| Pn-0 | 7307 | France |
| Pna-17 | 7523 | United States of America |
| Pro-0 | 8213 | Spain |
| Pu2-23 | 6951 | Czech Republic |
| PUZ24 | — | — |
| Ra-0 | 6958 | France |
| Rak-2 | 8365 | Czech Republic |
| Ren-1 | 6959 | France |
| Rev-2 | 6076 | Sweden |
| Rhen-1 | 7316 | Netherlands |
| Rmx-A180 | 7525 | United States of America |
| ROM-1 | 267 | France |
| RRS | — | — |
| Rsch-4 | 8374 | Russia |
| S96 | 7472 | Netherlands |
| Sanna-2 | 8376 | Sweden |
| Sap-0 | 8378 | Czech Republic |
| Sapporo-0 | 7330 | Japan |
| Sav-0 | 7340 | Czech Republic |
| Sav-1 | — | — |
| Se-0 | 6961 | Spain |

| | | |
|---|---|---|
| Sg-1 | 7344 | Germany |
| Shahdara | 6962 | Tajikistan |
| Si-0 | 7337 | Germany |
| SLSP-30 | 2274 | United States of America |
| Sp-0 | 7343 | Germany |
| Sq-8 | 6967 | United Kingdom |
| St-0 | 8387 | Sweden |
| Ste-0 | 7346 | Germany |
| Ste-3 | 2290 | United States of America |
| T1040 | 6094 | Sweden |
| T1060 | 6096 | Sweden |
| T1080 | 6098 | Sweden |
| T1110 | 6100 | Sweden |
| T1130 | 6102 | Sweden |
| T540 | 6112 | Sweden |
| T690 | 6124 | Sweden |
| Tad01 | 6169 | Sweden |
| TDr-1 | 6188 | Sweden |
| TDr-18 | 6203 | Sweden |
| TDr-3 | 6190 | Sweden |
| TDr-8 | 6194 | Sweden |
| Tha-1 | 7353 | Netherlands |
| Tiv-1 | 7355 | Italy |
| Tomegap-2 | 6242 | Sweden |
| Tottarp-2 | 6243 | Sweden |
| TOU-A1-115 | 281 | France |
| TOU-A1-116 | 282 | France |
| TOU-A1-12 | — | France |
| TOU-A1-43 | 321 | France |
| TOU-A1-62 | 328 | France |
| TOU-A1-67 | — | France |
| TOU-A1-96 | 357 | France |
| TOU-C-3 | 362 | France |
| TOU-E-11 | 366 | France |
| TOU-H-12 | 373 | France |
| TOU-H-13 | 374 | France |
| TOU-I-17 | 378 | France |
| TOU-I-2 | 379 | France |
| TOU-I-6 | 380 | France |
| TOU-J-3 | 383 | France |
| TOU-K-3 | 386 | France |
| Ts-1 | 6970 | Spain |
| Tscha-1 | 7372 | Austria |
| Tsu-0 | 7373 | Japan |

| | | |
|---|---|---|
| Ty-0 | 7351 | United Kingdom |
| Udul1-34 | 6318 | Czech Republic |
| Uk-1 | 7378 | Germany |
| UKCW06202 | — | United Kingdom |
| UKID22 | 5729 | United Kingdom |
| UKID37 | 5742 | United Kingdom |
| UKID48 | 5753 | United Kingdom |
| UKNW06-059 | 5380 | United Kingdom |
| UKNW06-060 | 5381 | United Kingdom |
| UKNW06-386 | 5565 | United Kingdom |
| UKNW06-436 | 5606 | United Kingdom |
| UKNW06-460 | 5628 | United Kingdom |
| UKSE06-062 | 4997 | United Kingdom |
| UKSE06-192 | 5056 | United Kingdom |
| UKSE06-272 | 5116 | United Kingdom |
| UKSE06-349 | 5158 | United Kingdom |
| UKSE06-351 | 5160 | United Kingdom |
| UKSE06-429 | 5207 | United Kingdom |
| UKSE06-466 | 5232 | United Kingdom |
| UKSE06-482 | 5245 | United Kingdom |
| UKSE06-520 | 5264 | United Kingdom |
| UKSE06-628 | 5341 | United Kingdom |
| Ull2-3 | 6973 | Sweden |
| Ull2-5 | 6974 | Sweden |
| Ull3-4 | 6413 | Sweden |
| Uod-7 | 6976 | Austria |
| Utrecht | 7382 | Netherlands |
| Van-0 | 6977 | Canada |
| Var2-1 | 7516 | Sweden |
| VOU-1 | 390 | France |
| VOU-2 | 392 | France |
| Wa-1 | 7394 | Poland |
| Wag-3 | 7390 | Netherlands |
| Wag-4 | 7391 | Netherlands |
| Wag-5 | 7392 | Netherlands |
| Wc-2 | 7405 | Germany |
| Wei-0 | 6979 | Switzerland |
| Wil-1 | — | Lithuania |
| Wl-0 | 7411 | Germany |
| Ws | 7397 | Russia |
| Ws-0 | 6980 | Russia |
| Wt-3 | 7408 | Germany |
| Wt-5 | 6982 | Germany |
| Yo-0 | 6983 | United States of America |

| | | |
|---|---|---|
| Zdr-6 | 6985 | Czech Republic |
| Zdrl2-24 | 6448 | Czech Republic |
| Zdrl2-25 | 6449 | Czech Republic |
| Zu-1 | 7418 | Switzerland |

**Supplementary Table 2.** Descriptive statistics of the effective ($F_q'/F_m'$) and the maximal efficiency ($F_v/F_m$) of photosystem II, projected rosette area, and relative growth rate (RGR).

Means (± 95% confidence interval) are shown together with the minimum and the maximum values. RGR was calculated from the projected rosette area measured on two consecutive days. For example, RGR at 16 days after stratification (DAS) was calculated from the projected rosette area measured at 16 and 17 DAS.

| Trait | $T_{night}$ | DAS | Mean | Minimum | Maximum |
|---|---|---|---|---|---|
| $F_q'/F_m'$ | 15°C | 16 | 0.43±0.003 | 0.33 | 0.48 |
| | | 17 | 0.44±0.003 | 0.31 | 0.49 |
| | | 18 | 0.45±0.002 | 0.34 | 0.50 |
| | 20°C | 16 | 0.45±0.002 | 0.33 | 0.49 |
| | | 17 | 0.46±0.002 | 0.36 | 0.50 |
| | | 18 | 0.47±0.002 | 0.38 | 0.51 |
| $F_v/F_m$ | 15°C | 16 | 0.77±0.001 | 0.69 | 0.79 |
| | | 17 | 0.77±0.001 | 0.65 | 0.80 |
| | | 18 | 0.78±0.001 | 0.71 | 0.80 |
| | 20°C | 16 | 0.76±0.001 | 0.72 | 0.78 |
| | | 17 | 0.77±0.001 | 0.69 | 0.79 |
| | | 18 | 0.77±0.001 | 0.71 | 0.79 |
| Projected rosette area ($cm^2$) | 15°C | 16 | 0.59±0.02 | 0.25 | 1.07 |
| | | 17 | 0.85±0.02 | 0.15 | 1.60 |
| | | 18 | 1.25±0.03 | 0.23 | 2.39 |
| | 20°C | 16 | 1.20±0.03 | 0.68 | 2.35 |
| | | 17 | 1.76±0.04 | 0.83 | 3.27 |
| | | 18 | 2.54±0.06 | 1.20 | 4.87 |
| RGR (% $d^{-1}$) | 15°C | 16 | 38.3±0.58 | 23.1 | 50.6 |
| | | 17 | 38.5±0.47 | 24.5 | 50.9 |
| | 20°C | 16 | 38.9±0.38 | 30.4 | 48.2 |
| | | 17 | 37.4±0.34 | 28.0 | 44.7 |

**Supplementary Table 3.** Single nucleotide polymorphisms (SNPs) associated with long-term response of $F_q'/F_m'$ to night temperature.

The SNP-trait associations were considered significant when $-\log_{10}(P) \geq$ FDR (false discovery rate) threshold (0.05) and/or $-\log_{10}(P) > 4.0$ on at least two days. No. refers to the SNP number shown in Figure 2. Allele 1/2 shows the major/minor allele. The allele effects were estimated for the minor alleles and given in the unit of $F_q'/F_m'$. The $r^2$ values correspond to the proportions of the variance explained by individual SNPs or sets of SNPs (simultaneous fit). For pairs of SNPs that were in linkage disequilibrium (underlined), only one of them (in parentheses) was included in the simultaneous fit. DAS, days after stratification; MAF, minor allele frequency. *SNPs are located in intergenic regions.

| No. | SNP | $T_{night}$ | DAS | Allele 1/2 | Col-0 allele | MAF (%) | $-\log_{10}$ ($P$) | Effect | $r^2$ (%) | Locus | Gene annotation |
|---|---|---|---|---|---|---|---|---|---|---|---|
| 1 | <u>Chr2_1138854</u> | 15°C | 16 | C/T | C | 12.33 | 4.63 | -0.009 | 8.77 | AT2G03730 | ACT DOMAIN REPEAT 5 |
| 2 | <u>(Chr2_1138985)</u> | 15°C | 16 | C/T | C | 11.99 | 5.14 | -0.010 | 9.73 | AT2G03730 | ACT DOMAIN REPEAT 5 |
| 3 | Chr3_6651631 | 15°C | 16 | A/C | A | 21.23 | 5.80 | -0.008 | 9.64 | * | * |
| 4 | Chr3_16153315 | 15°C | 16 | T/C | T | 45.89 | 4.10 | 0.006 | 6.81 | AT3G44560 | FATTY ACID REDUCTASE 8 |
| | | Simultaneous fit | | | | | | | 18.30 | | |
| 5 | Chr1_846217 | 15°C | 17 | A/C | C | 41.30 | 4.44 | 0.006 | 10.02 | AT1G03410 | 2-oxoglutarate and Fe (II)-dependent oxygenase superfamily protein |
| 6 | Chr1_10444149 | 15°C | 17 | A/G | A | 24.23 | 4.91 | -0.007 | 7.02 | AT1G29830 | magnesium transporter CorA-like family protein |
| 2 | Chr2_1138985 | 15°C | 17 | C/T | C | 11.95 | 4.07 | -0.008 | 8.40 | AT2G03730 | ACT DOMAIN REPEAT 5 |
| 7 | Chr3_1409314 | 15°C | 17 | T/C | T | 2.39 | 6.54 | -0.022 | 12.43 | AT3G05050 | protein kinase superfamily protein |
| 3 | Chr3_6651631 | 15°C | 17 | A/C | A | 21.16 | 4.35 | -0.007 | 7.97 | * | * |
| 8 | <u>Chr3_9768774</u> | 15°C | 17 | A/C | A | 6.48 | 4.60 | -0.011 | 8.24 | * | * |
| 9 | <u>(Chr3_9774541)</u> | 15°C | 17 | A/G | A | 6.48 | 4.60 | -0.011 | 8.18 | * | * |
| 4 | Chr3_16153315 | 15°C | 17 | T/C | T | 45.73 | 4.15 | 0.005 | 6.61 | AT3G44560 | FATTY ACID REDUCTASE 8 |
| 10 | Chr4_7788139 | 15°C | 17 | C/T | C | 12.97 | 4.88 | -0.009 | 9.93 | AT4G13400 | 2-oxoglutarate and Fe (II)-dependent oxygenase superfamily protein |

| | | | | | | | | | | | |
|---|---|---|---|---|---|---|---|---|---|---|---|
| 11 | Chr4_7788811 | 15°C | 17 | A/T | A | 5.46 | 4.77 | -0.012 | 9.21 | AT4G13400 | 2-oxoglutarate and Fe (II)-dependent oxygenase superfamily protein |
| 12 | Chr5_3676362 | 15°C | 17 | T/A | T | 13.99 | 4.64 | -0.008 | 9.17 | AT5G11490 | adaptin family protein |
| 13 | Chr5_5649958 | 15°C | 17 | C/T | T | 29.35 | 4.19 | -0.006 | 4.97 | AT5G17170 | rubredoxin family protein |
| | | Simultaneous fit | | | | | | | 32.01 | | |
| 5 | Chr1_846217 | 15°C | 18 | A/C | C | 41.30 | 4.40 | 0.006 | 9.96 | AT1G03410 | 2-oxoglutarate and Fe (II)-dependent oxygenase superfamily protein |
| 6 | Chr1_10444149 | 15°C | 18 | A/G | A | 24.23 | 4.69 | -0.006 | 6.59 | AT1G29830 | magnesium transporter CorA-like family protein |
| 1 | Chr2_1138854 | 15°C | 18 | C/T | C | 12.29 | 4.11 | -0.008 | 7.87 | AT2G03730 | ACT DOMAIN REPEAT 5 |
| 2 | (Chr2_1138985) | 15°C | 18 | C/T | C | 11.95 | 4.69 | -0.008 | 9.16 | AT2G03730 | ACT DOMAIN REPEAT 5 |
| 7 | Chr3_1409314 | 15°C | 18 | T/C | T | 2.39 | 5.21 | -0.018 | 9.93 | AT3G05050 | protein kinase superfamily protein |
| 8 | Chr3_9768774 | 15°C | 18 | A/C | A | 6.48 | 4.30 | -0.010 | 7.83 | * | * |
| 9 | (Chr3_9774541) | 15°C | 18 | A/G | A | 6.48 | 4.30 | -0.010 | 7.58 | * | * |
| 10 | Chr4_7788139 | 15°C | 18 | C/T | C | 12.97 | 5.02 | -0.008 | 9.74 | AT4G13400 | 2-oxoglutarate and Fe (II)-dependent oxygenase superfamily protein |
| 11 | Chr4_7788811 | 15°C | 18 | A/T | A | 5.46 | 4.37 | -0.011 | 7.90 | AT4G13400 | 2-oxoglutarate and Fe (II)-dependent oxygenase superfamily protein |
| 12 | Chr5_3676362 | 15°C | 18 | T/A | T | 13.99 | 5.09 | -0.008 | 10.28 | AT5G11490 | adaptin family protein |
| 13 | Chr5_5649958 | 15°C | 18 | C/T | T | 29.35 | 4.21 | -0.006 | 5.01 | AT5G17170 | rubredoxin family protein |
| | | Simultaneous fit | | | | | | | 29.09 | | |
| 14 | Chr1_13939977 | 20°C | 16 | A/G | G | 31.85 | 7.88 | -0.004 | 6.63 | * | * |
| 2 | Chr2_1138985 | 20°C | 16 | C/T | C | 11.99 | 8.54 | -0.007 | 11.37 | AT2G03730 | ACT DOMAIN REPEAT 5 |
| 15 | Chr2_11170979 | 20°C | 16 | A/G | A | 2.05 | 9.63 | -0.015 | 18.17 | AT2G26250 | 3-KETOACYL-COA SYNTHASE 10 |

| | | | | | | | | | | | |
|---|---|---|---|---|---|---|---|---|---|---|---|
| 16 | Chr3_3946441 | 20°C | 16 | C/T | C | 16.78 | 5.40 | -0.004 | 9.00 | AT3G12410 | polynucleotidyl transferase |
| 17 | Chr4_7156570 | 20°C | 16 | A/T | T | 11.30 | 7.98 | -0.006 | 2.62 | AT4G11910 | STAY-GREEN-like protein |
| 18 | Chr4_7653545 | 20°C | 16 | G/T | G | 31.16 | 7.54 | 0.004 | 3.82 | * | * |
| 19 | Chr4_8562879 | 20°C | 16 | T/G | T | 13.36 | 6.53 | -0.005 | 9.25 | AT4G14980 | cysteine/histidine-rich C1 domain family protein |
| 20 | Chr5_14578542 | 20°C | 16 | C/A | C | 41.44 | 6.91 | -0.004 | 6.49 | AT5G36935 | transposable element |
| 21 | Chr5_15330192 | 20°C | 16 | G/T | T | 41.44 | 7.05 | 0.004 | 4.22 | AT5G38350 | disease resistance protein (NBS-LRR class) family |
| | | Simultaneous fit | | | | | | | 39.87 | | |
| 22 | Chr1_2029747 | 20°C | 17 | A/T | T | 47.78 | 9.00 | -0.004 | 4.88 | AT1G06630 | F-box/RNI-like superfamily protein |
| 23 | Chr1_8764728 | 20°C | 17 | C/T | C | 44.71 | 6.76 | -0.003 | 3.01 | * | * |
| 15 | Chr2_11170979 | 20°C | 17 | A/G | A | 2.05 | 23.59 | -0.024 | 19.09 | AT2G26250 | 3-KETOACYL-COA SYNTHASE 10 |
| 16 | Chr3_3946441 | 20°C | 17 | C/T | C | 16.72 | 7.06 | -0.004 | 10.05 | AT3G12410 | polynucleotidyl transferase |
| 24 | Chr4_2580731 | 20°C | 17 | G/A | A | 30.38 | 6.07 | -0.003 | 1.62 | AT4G05040 | ankyrin repeat family protein |
| 25 | Chr4_11273457 | 20°C | 17 | G/A | G | 5.80 | 8.95 | -0.008 | 8.68 | AT4G21120 | AMINO ACID TRANSPORTER 1 |
| 26 | Chr4_11827105 | 20°C | 17 | T/C | T | 23.21 | 7.19 | 0.004 | 3.62 | * | * |
| 20 | Chr5_14578542 | 20°C | 17 | C/A | C | 41.30 | 8.57 | -0.004 | 6.50 | AT5G36935 | transposable element |
| 27 | Chr5_18289242 | 20°C | 17 | A/G | G | 24.57 | 9.49 | -0.004 | 7.02 | AT5G45200 | disease resistance protein (TIR-NBS-LRR class) family |
| | | Simultaneous fit | | | | | | | 39.91 | | |
| 28 | Chr1_18957084 | 20°C | 18 | G/A | G | 6.14 | 7.25 | -0.007 | 11.19 | * | * |
| 29 | Chr2_8145104 | 20°C | 18 | A/G | G | 21.16 | 6.01 | 0.003 | 9.16 | AT2G18800 | XYLOGLUCAN ENDOTRANSGLUCOSYLASE/ HYDROLASE 21 |
| 15 | Chr2_11170979 | 20°C | 18 | A/G | A | 2.05 | 9.92 | -0.015 | 17.75 | AT2G26250 | 3-KETOACYL-COA SYNTHASE 10 |

| | | | | | | | | | | | |
|---|---|---|---|---|---|---|---|---|---|---|---|
| 30 | Chr3_3218698 | 20°C | 18 | G/A | G | 5.46 | 6.21 | -0.006 | 2.02 | AT3G10370 | FAD-dependent oxidoreductase family protein |
| 31 | Chr3_9123175 | 20°C | 18 | G/C | G | 20.14 | 8.57 | -0.004 | 6.71 | AT3G25030 | RING/U-box superfamily protein |
| 32 | Chr4_7274303 | 20°C | 18 | G/T | T | 38.91 | 11.24 | -0.004 | 8.14 | AT4G12170 | Thioredoxin superfamily protein |
| 33 | Chr4_11273410 | 20°C | 18 | G/C | G | 6.14 | 10.41 | -0.008 | 9.29 | AT4G21120 | AMINO ACID TRANSPORTER 1 |
| 34 | Chr5_9769302 | 20°C | 18 | T/C | T | 40.27 | 7.50 | -0.003 | 2.90 | AT5G27610 | ALWAYS EARLY 1 |
| 35 | Chr5_16900781 | 20°C | 18 | G/T | G | 46.08 | 6.49 | -0.003 | 1.86 | * | * |
| 27 | Chr5_18289242 | 20°C | 18 | A/G | G | 24.57 | 12.89 | -0.005 | 8.63 | AT5G45200 | disease resistance protein (TIR-NBS-LRR class) family |
| | | | Simultaneous fit | | | | | | 45.58 | | |

**Supplementary Table 4.** Number of single nucleotide polymorphisms (SNPs) used as predictors of $F_q'/F_m'$.

| $T_{night}$ | DAS | $-\log_{10}(P) \geq$ FDR and/or $-\log_{10}(P) > 4.0$ on at least two days | $-\log_{10}(P) \geq 3.0$ on the respective day |
|---|---|---|---|
| 15°C | 16 | 4 | 192 |
| | 17 | 12 | 231 |
| | 18 | 11 | 232 |
| 20°C | 16 | 9 | 192 |
| | 17 | 9 | 160 |
| | 18 | 10 | 260 |

DAS, days after stratification; FDR, false discovery rate; $T_{night}$, nighttime temperature.

**Supplementary Table 5.** Performance of gBLUP and Bayesian approaches in predicting $F_q'/F_m'$ for the 293 Arabidopsis accessions in the GWAS panel.

$F_q'/F_m'$ data of the 293 accessions were used to train prediction models based on 211,771 SNPs. Only at 16 days after stratification (DAS), one accession (T1080) was excluded. Predictive performance was assessed using root mean squared error (RMSE) and Pearson correlation coefficient (COR) averaged across three testing sets (± SE**)**.

| $T_{night}$ | Method | DAS | RMSE | COR |
|---|---|---|---|---|
| 15°C | gBLUP | 16 | 0.022±0.000 | 0.20±0.13 |
| | BayesA | 16 | 0.022±0.000 | 0.19±0.13 |
| | BayesB | 16 | 0.022±0.000 | 0.19±0.12 |
| | gBLUP | 17 | 0.022±0.002 | 0.23±0.06 |
| | BayesA | 17 | 0.022±0.002 | 0.24±0.06 |
| | BayesB | 17 | 0.022±0.002 | 0.23±0.06 |
| | gBLUP | 18 | 0.021±0.002 | 0.23±0.02 |
| | BayesA | 18 | 0.021±0.002 | 0.23±0.01 |
| | BayesB | 18 | 0.021±0.002 | 0.23±0.01 |
| 20°C | gBLUP | 16 | 0.022±0.001 | 0.15±0.09 |
| | BayesA | 16 | 0.022±0.001 | 0.17±0.08 |
| | BayesB | 16 | 0.022±0.001 | 0.15±0.07 |
| | gBLUP | 17 | 0.018±0.001 | 0.25±0.01 |
| | BayesA | 17 | 0.018±0.001 | 0.26±0.01 |
| | BayesB | 17 | 0.018±0.001 | 0.27±0.01 |
| | gBLUP | 18 | 0.016±0.000 | 0.21±0.03 |
| | BayesA | 18 | 0.016±0.001 | 0.23±0.02 |
| | BayesB | 18 | 0.016±0.001 | 0.23±0.02 |

gBLUP, genomic best linear unbiased prediction; $T_{night}$, night temperature.

**Supplementary Table 6.** 47 SNPs identified by GWAS with $-\log_{10}(P) \geq 3.0$ on three consecutive days in 15°C $T_{night}$.

Shown are the associations of 47 SNPs with $F_q'/F_m'$ at 16 and 17 days after stratification (DAS). The corresponding data for 18 DAS can be found in Table 1 together with locus and gene annotations. Allele 1/2 shows the major/minor allele. The allele effects were estimated for the minor alleles and given in the unit of $F_q'/F_m'$. The $r^2$ values give the proportion of the variance explained by individual SNPs or sets of SNPs (simultaneous fit). For pairs of SNPs that were in linkage disequilibrium (underlined), only one of them (in parentheses) was included in the simultaneous fit. MAF, minor allele frequency.

| SNP | DAS | Allele 1/2 | Col-0 allele | MAF (%) | $-\log_{10}(P)$ | Effect | $r^2$ (%) |
|---|---|---|---|---|---|---|---|
| Chr1_846217 | 16 | A/C | C | 41.44 | 4.00 | 0.006 | 9.65 |
| Chr1_873447 | 16 | A/T | A | 34.25 | 3.41 | -0.006 | 8.45 |
| Chr1_884146 | 16 | G/A | G | 37.33 | 3.26 | -0.005 | 7.24 |
| Chr1_6478834 | 16 | A/T | A | 23.97 | 3.65 | 0.007 | 3.08 |
| Chr1_10027869 | 16 | T/C | T | 9.25 | 3.56 | 0.009 | 7.27 |
| Chr1_19515673 | 16 | C/G | C | 38.01 | 3.08 | 0.005 | 7.04 |
| Chr1_23436110 | 16 | G/C | G | 6.85 | 3.58 | -0.010 | 5.75 |
| Chr2_1138854 | 16 | C/T | C | 12.33 | 4.63 | -0.009 | 9.19 |
| Chr2_1138985 | 16 | C/T | C | 11.99 | 5.14 | -0.010 | 10.08 |
| Chr2_1139010 | 16 | A/C | A | 14.73 | 4.46 | -0.008 | 8.51 |
| Chr2_2884161 | 16 | C/T | T | 31.16 | 3.01 | -0.005 | 2.93 |
| Chr2_8145104 | 16 | A/G | G | 21.23 | 3.62 | 0.007 | 7.17 |
| Chr2_11170979 | 16 | A/G | A | 2.05 | 3.77 | -0.019 | 8.72 |
| Chr2_13684149 | 16 | G/A | G | 13.70 | 3.46 | -0.007 | 6.67 |
| Chr2_14059450 | 16 | C/A | C | 29.11 | 3.28 | -0.006 | 6.99 |
| Chr3_1275424 | 16 | C/T | C | 16.78 | 3.47 | 0.007 | 3.59 |
| Chr3_3395109 | 16 | C/G | C | 9.59 | 3.18 | 0.009 | 6.94 |

| | | | | | | | |
|---|---|---|---|---|---|---|---|
| Chr3_6651631 | 16 | A/C | A | 21.23 | 5.80 | -0.008 | 10.12 |
| Chr3_9636722 | 16 | T/G | T | 34.93 | 4.27 | -0.006 | 8.09 |
| Chr3_10582418 | 16 | C/T | T | 47.60 | 3.66 | 0.006 | 2.76 |
| Chr3_10874763 | 16 | T/A | A | 39.73 | 3.20 | -0.005 | 3.45 |
| Chr3_10898637 | 16 | G/A | G | 28.77 | 4.11 | -0.006 | 9.02 |
| Chr3_16153315 | 16 | T/C | T | 45.89 | 4.10 | 0.006 | 6.70 |
| Chr3_19054365 | 16 | A/G | A | 3.08 | 3.91 | -0.016 | 8.34 |
| Chr3_19257874 | 16 | T/C | T | 9.59 | 3.41 | 0.009 | 6.43 |
| Chr3_21427556 | 16 | G/A | G | 12.67 | 4.89 | -0.009 | 8.74 |
| Chr3_21432797 | 16 | C/T | C | 21.23 | 4.87 | -0.008 | 8.25 |
| Chr3_21432851 | 16 | C/T | C | 21.92 | 4.62 | -0.007 | 8.04 |
| Chr3_22377074 | 16 | A/G | A | 36.64 | 3.70 | -0.005 | 6.10 |
| Chr4_5411416 | 16 | C/G | C | 19.18 | 3.22 | -0.006 | 6.09 |
| Chr4_5633114 | 16 | C/A | C | 3.08 | 4.30 | -0.016 | 8.88 |
| Chr4_7788139 | 16 | C/T | C | 12.67 | 3.43 | -0.008 | 8.00 |
| Chr4_14017514 | 16 | C/T | C | 15.07 | 3.48 | -0.007 | 6.66 |
| Chr4_14030315 | 16 | T/G | T | 16.10 | 3.10 | -0.006 | 5.89 |
| Chr4_15903731 | 16 | C/G | C | 2.05 | 3.75 | -0.019 | 8.28 |
| Chr4_18442418 | 16 | A/G | A | 12.67 | 3.90 | -0.008 | 7.10 |
| Chr5_700043 | 16 | A/G | A | 17.12 | 4.13 | -0.008 | 5.20 |
| Chr5_3284989 | 16 | C/A | A | 40.75 | 3.01 | 0.005 | 3.78 |
| Chr5_5536626 | 16 | A/C | A | 19.18 | 3.11 | -0.006 | 7.55 |
| Chr5_5538353 | 16 | G/A | G | 18.15 | 3.22 | -0.007 | 7.46 |
| Chr5_5620806 | 16 | C/T | T | 46.58 | 3.99 | -0.005 | 7.18 |
| Chr5_5627597 | 16 | T/G | T | 29.79 | 3.24 | -0.005 | 6.07 |
| Chr5_5627893 | 16 | A/G | G | 35.62 | 3.38 | -0.005 | 4.96 |
| Chr5_9760768 | 16 | C/A | C | 6.16 | 3.74 | -0.011 | 7.38 |
| Chr5_18289242 | 16 | A/G | G | 24.32 | 3.21 | -0.006 | 6.29 |

| | | | | | | | |
|---|---|---|---|---|---|---|---|
| Chr5_21767983 | 16 | T/C | T | 3.08 | 3.93 | -0.016 | 9.22 |
| Chr5_23908100 | 16 | C/A | A | 46.58 | 3.04 | 0.005 | 6.64 |
| | | | Simultaneous fit | | | | 49.09 |
| Chr1_846217 | 17 | A/C | C | 41.30 | 4.44 | 0.006 | 10.25 |
| Chr1_873447 | 17 | A/T | A | 34.47 | 3.79 | -0.006 | 8.87 |
| Chr1_884146 | 17 | G/A | G | 37.54 | 3.80 | -0.005 | 7.32 |
| Chr1_6478834 | 17 | A/T | A | 23.89 | 3.25 | 0.006 | 2.95 |
| Chr1_10027869 | 17 | T/C | T | 9.22 | 3.09 | 0.008 | 6.43 |
| Chr1_19515673 | 17 | C/G | C | 37.88 | 3.29 | 0.005 | 7.37 |
| Chr1_23436110 | 17 | G/C | G | 6.83 | 3.01 | -0.009 | 5.05 |
| Chr2_1138854 | 17 | C/T | C | 12.29 | 3.67 | -0.008 | 7.62 |
| Chr2_1138985 | 17 | C/T | C | 11.95 | 4.07 | -0.008 | 8.25 |
| Chr2_1139010 | 17 | A/C | A | 14.68 | 3.35 | -0.007 | 6.25 |
| Chr2_2884161 | 17 | C/T | T | 31.40 | 3.56 | -0.005 | 3.68 |
| Chr2_8145104 | 17 | A/G | G | 21.16 | 4.05 | 0.007 | 8.09 |
| Chr2_11170979 | 17 | A/G | A | 2.05 | 3.81 | -0.018 | 8.14 |
| Chr2_13684149 | 17 | G/A | G | 13.65 | 3.09 | -0.006 | 6.02 |
| Chr2_14059450 | 17 | C/A | C | 29.01 | 3.19 | -0.005 | 6.72 |
| Chr3_1275424 | 17 | C/T | C | 16.72 | 3.44 | 0.007 | 3.52 |
| Chr3_3395109 | 17 | C/G | C | 9.56 | 3.05 | 0.008 | 7.12 |
| Chr3_6651631 | 17 | A/C | A | 21.16 | 4.35 | -0.007 | 8.13 |
| Chr3_9636722 | 17 | T/G | T | 34.81 | 3.52 | -0.005 | 7.25 |
| Chr3_10582418 | 17 | C/T | T | 47.44 | 3.53 | 0.005 | 2.80 |
| Chr3_10874763 | 17 | T/A | A | 39.93 | 3.86 | -0.005 | 4.21 |
| Chr3_10898637 | 17 | G/A | G | 28.67 | 3.29 | -0.005 | 7.78 |
| Chr3_16153315 | 17 | T/C | T | 45.73 | 4.15 | 0.005 | 6.59 |
| Chr3_19054365 | 17 | A/G | A | 3.07 | 4.03 | -0.015 | 9.16 |
| Chr3_19257874 | 17 | T/C | T | 9.56 | 3.23 | 0.008 | 5.63 |

| | | | | | | | |
|---|---|---|---|---|---|---|---|
| Chr3_21427556 | 17 | G/A | G | 12.63 | 3.82 | -0.008 | 7.72 |
| Chr3_21432797 | 17 | C/T | C | 21.16 | 3.41 | -0.006 | 6.72 |
| Chr3_21432851 | 17 | C/T | C | 21.84 | 3.28 | -0.006 | 6.38 |
| Chr3_22377074 | 17 | A/G | A | 36.86 | 3.43 | -0.005 | 5.53 |
| Chr4_5411416 | 17 | C/G | C | 19.11 | 3.25 | -0.006 | 6.72 |
| Chr4_5633114 | 17 | C/A | C | 3.07 | 3.63 | -0.014 | 7.79 |
| Chr4_7788139 | 17 | C/T | C | 12.97 | 4.88 | -0.009 | 10.06 |
| Chr4_14017514 | 17 | C/T | C | 15.36 | 4.03 | -0.007 | 7.78 |
| Chr4_14030315 | 17 | T/G | T | 16.38 | 3.57 | -0.007 | 6.79 |
| Chr4_15903731 | 17 | C/G | C | 2.05 | 3.96 | -0.018 | 9.24 |
| Chr4_18442418 | 17 | A/G | A | 12.63 | 3.26 | -0.007 | 6.02 |
| Chr5_700043 | 17 | A/G | A | 17.06 | 3.45 | -0.006 | 4.43 |
| Chr5_3284989 | 17 | C/A | A | 40.61 | 3.43 | 0.005 | 4.47 |
| Chr5_5536626 | 17 | A/C | A | 19.11 | 3.25 | -0.006 | 8.25 |
| Chr5_5538353 | 17 | G/A | G | 18.09 | 3.10 | -0.006 | 7.44 |
| Chr5_5620806 | 17 | C/T | T | 46.76 | 4.60 | -0.006 | 8.00 |
| Chr5_5627597 | 17 | T/G | T | 30.03 | 3.53 | -0.005 | 6.15 |
| Chr5_5627893 | 17 | A/G | G | 35.84 | 3.51 | -0.005 | 5.48 |
| Chr5_9760768 | 17 | C/A | C | 6.14 | 4.58 | -0.012 | 8.28 |
| Chr5_18289242 | 17 | A/G | G | 24.57 | 3.58 | -0.006 | 6.76 |
| Chr5_21767983 | 17 | T/C | T | 3.07 | 3.57 | -0.014 | 8.87 |
| Chr5_23908100 | 17 | C/A | A | 46.42 | 3.59 | 0.005 | 7.12 |
| | | | Simultaneous fit | | | | 47.08 |

**Supplementary Table 7.** Performance of gBLUP in predicting $F_q'/F_m'$ for the Arabidopsis GWAS panel in 15°C $T_{night}$ based on four different sets of SNPs.

Predictors were 211,771 SNPs or the GWAS-derived SNPs with $-\log_{10}$ (*P*) ≥ FDR threshold (0.05) and/or $-\log_{10}$ (*P*) > 4.0 on at least two days (a), $-\log_{10}$ (*P*) ≥ 3.0 on the respective day (b), and 47 SNPs that satisfied $-\log_{10}$ (*P*) ≥ 3.0 on all three days (c). Predictive performance was assessed using root mean squared error (RMSE) and Pearson correlation coefficient (COR) averaged across three testing sets (± SE).

| DAS | 211,771 SNPs | | FDR threshold[a] | | -log10 (*P*) ≥ 3.0[b] | | 47 SNPs[c] | |
|---|---|---|---|---|---|---|---|---|
| | RMSE | COR | RMSE | COR | RMSE | COR | RMSE | COR |
| 16 | 0.022±0.001 | 0.20±0.13 | 0.021±0.001 | 0.32±0.06 | 0.014±0.001 | 0.79±0.01 | 0.016±0.002 | 0.71±0.05 |
| 17 | 0.022±0.002 | 0.23±0.06 | 0.020±0.001 | 0.48±0.08 | 0.012±0.000 | 0.84±0.04 | 0.017±0.002 | 0.69±0.04 |
| 18 | 0.021±0.002 | 0.23±0.02 | 0.018±0.000 | 0.51±0.12 | 0.012±0.001 | 0.82±0.06 | 0.016±0.001 | 0.67±0.03 |

DAS, days after stratification; FDR, false discovery rate; $T_{night}$, night temperature.

**Supplementary Table 8.** Performance of gBLUP in predicting $F_q'/F_m'$ for the Arabidopsis GWAS panel in 20°C night temperature ($T_{night}$) at 18 days after stratification (DAS) based on SNPs identified in the same $T_{night}$ but on wrong days, or on the same day but in 15°C $T_{night}$.

For comparison, predictive performance based on the corresponding set of SNPs (*identified in 20°C $T_{night}$ at 18 DAS) is also shown. Predictive performance was assessed using root mean squared error (RMSE) and Pearson correlation coefficient (COR) averaged across three test sets (± SE). The associations of all SNPs had $-\log_{10}(P) \geq 3.0$.

| $T_{night}$ | DAS | RMSE | COR |
|---|---|---|---|
| 15°C | 18 | 0.013±0.000 | 0.62±0.06 |
| 20°C | 16 | 0.012±0.000 | 0.67±0.05 |
| 20°C | 17 | 0.010±0.000 | 0.78±0.05 |
| *20°C | 18 | 0.008±0.001 | 0.86±0.03 |

**Supplementary Table 9.** Performance of XGBoost in predicting $F_q'/F_m'$ based on the genotype data adjusted and non-adjusted for population structure.

Prediction was based on 211,771 SNPs. Genotypes were adjusted for population structure using 250 principal components. Validation of the XGBoost model in the test set was performed using parameters selected in a grid search (see Supplementary Table 11 for hyperparameters evaluated in the grid search). Predictive performance was assessed by root mean squared error (RMSE ±SE) averaged across three test sets.

| $T_{night}$ | DAS | RMSE | |
|---|---|---|---|
| | | Non-adjusted | Adjusted |
| 15°C | 16 | 0.023±0.002 | 0.023±0.002 |
| | 17 | 0.023±0.003 | 0.022±0.003 |
| | 18 | 0.021±0.002 | 0.021±0.002 |
| 20°C | 16 | 0.022±0.001 | 0.022±0.001 |
| | 17 | 0.017±0.002 | 0.018±0.001 |
| | 18 | 0.016±0.001 | 0.016±0.001 |

DAS, days after stratification; $T_{night}$, night temperature.

**Supplementary Table 10.** List of nine accessions in the validation experiment.

Ecotype ID refers to the accession identification in the genotype data. Alongside these accessions, Col-0 was included as reference in the validation experiment.

| Accession | Ecotype ID | Country |
|---|---|---|
| CUR-8 | 86 | France |
| PAR-8 | 262 | France |
| Paw-4 | 2151 | United States of America |
| Kyl-1 | 5751 | United Kingdom |
| DraIV3-8 | 5922 | Czech Republic |
| Ei-2 | 6915 | Germany |
| LL-0 | 6933 | Spain |
| Bs-1 | 8270 | Switzerland |
| Sim-1 | 9442 | Sweden |

**Supplementary Table 11.** Hyperparameters evaluated in the grid search to maximize the predictive performance of XGBoost.

| Hyperparameter | Values |
|---|---|
| Maximal depth of a tree | 3, 5, 6, 10 |
| Learning rate | 0.0005, 0.005, 0.05, 0.01 |
| Subsampling of training set | 0.8, 0.9 |
| Number of estimators | 100, 500 |

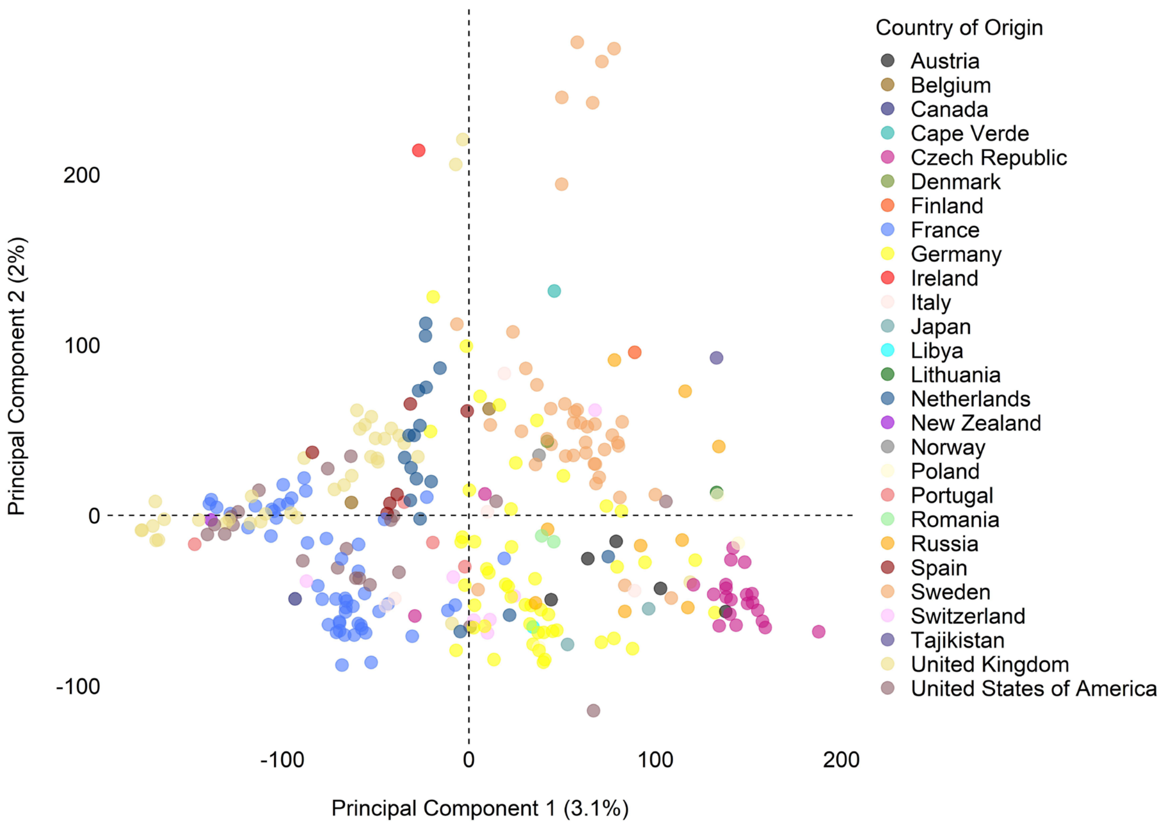


**Supplementary Fig. 1:** Population structure of the 293 Arabidopsis accessions. Principal component (PC) analysis was based on the 211,771 SNPs. Percentages of the total variance explained by PC1 and PC2 are 3.1% and 2%, respectively. Circles represent individual accessions (color-coded according to their country of origin).

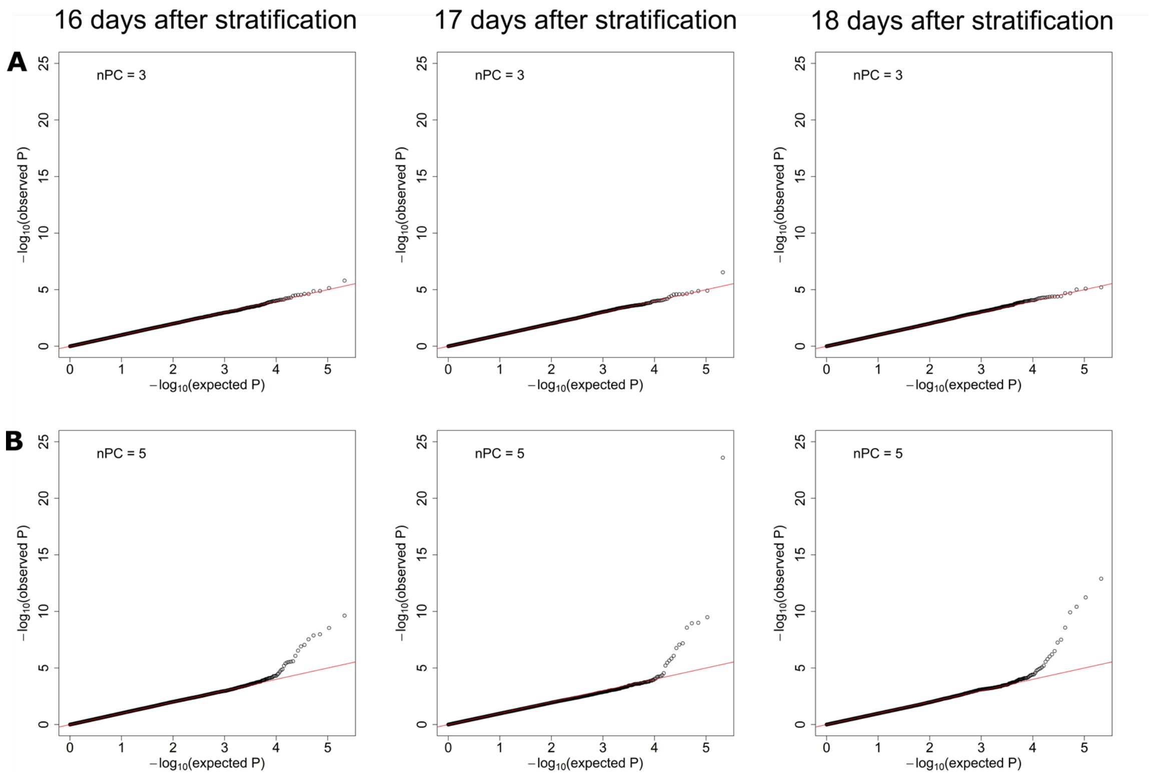


**Supplementary Fig. 2:** Q-Q plots of the observed and the expected *P* values distribution from the GWAS on $F_q'/F_m'$ in the 293 accessions at 16, 17 and 18 days after stratification. A, 15°C $T_{night}$. B, 20°C $T_{night}$. nPC indicates the number of principal components included as covariates in FarmCPU.

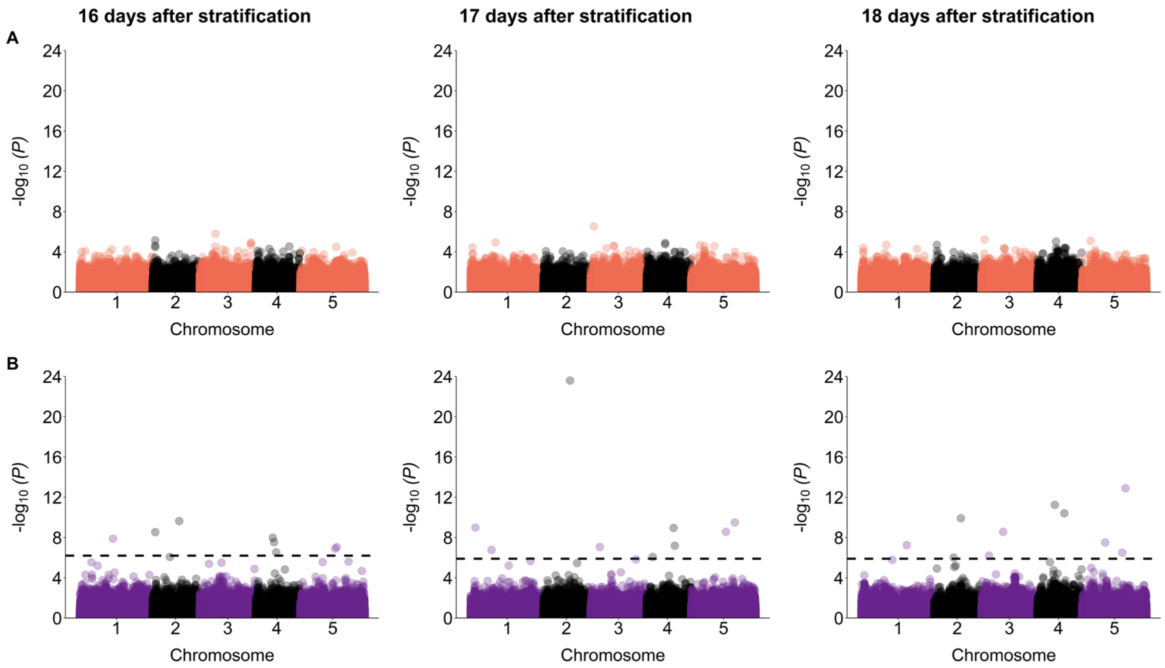


**Supplementary Fig. 3.** Manhattan plots of the GWAS on $F_q'/F_m'$ in the 293 accessions. A, 15°C $T_{night}$. B, 20°C $T_{night}$. Dashed lines in (B) show the false discovery rate threshold (0.05).

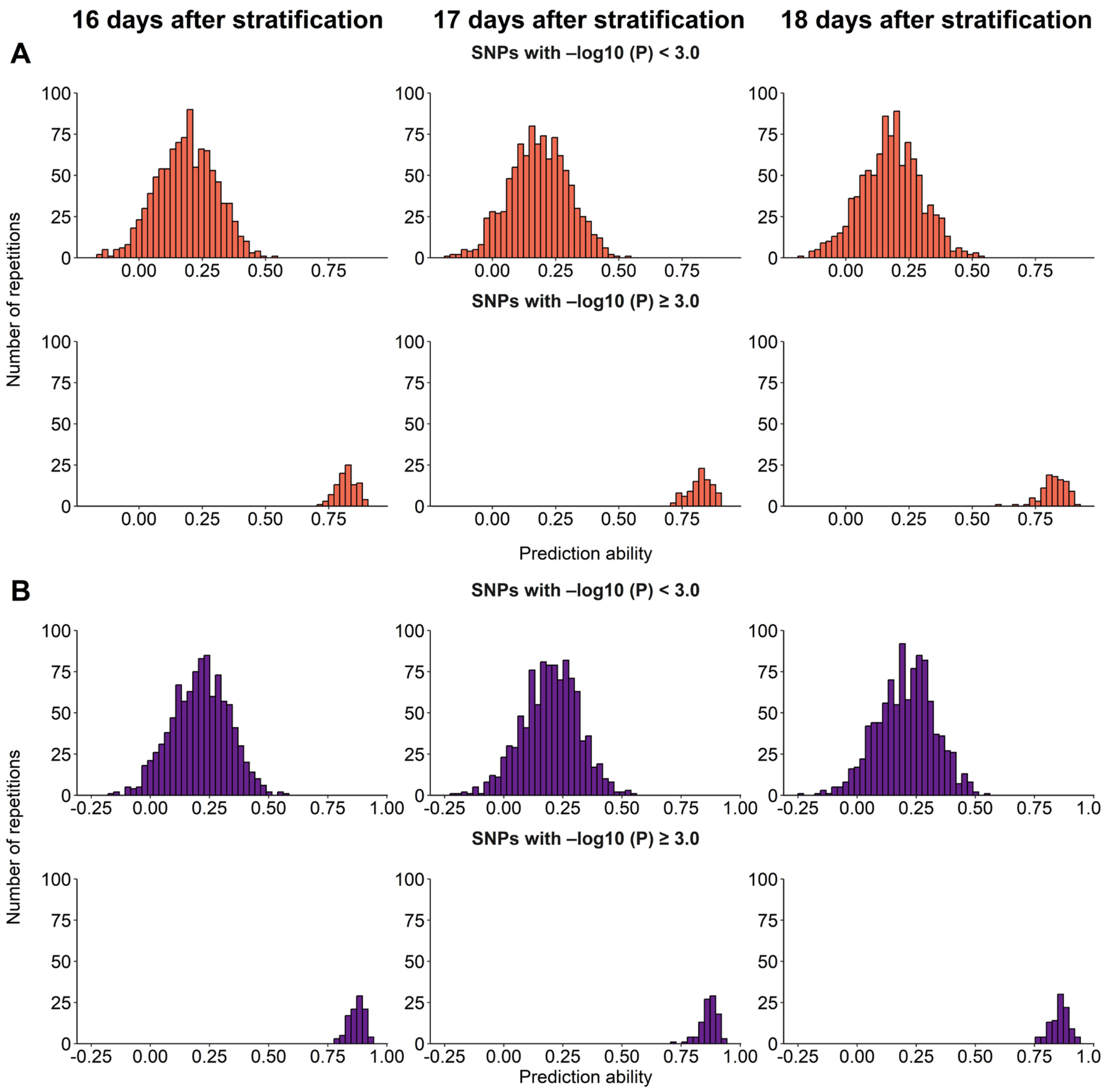


**Supplementary Fig. 4:** Distribution of prediction ability of gBLUP models for $F_q'/F_m'$ based on the GWAS-derived SNPs with $–\log_{10}$ ($P$) < 3 or ≥ 3. A, 15°C $T_{night}$. B, 20°C $T_{night}$. For the plots of $-\log_{10}$ ($P$) < 3, prediction ability was calculated by randomly selecting the same number of SNPs as for $-\log_{10}$ ($P$) ≥ 3. Given the large number of SNPs with $-\log_{10}$ ($P$) < 3, prediction ability was calculated as the median across 992-1,000 replications.

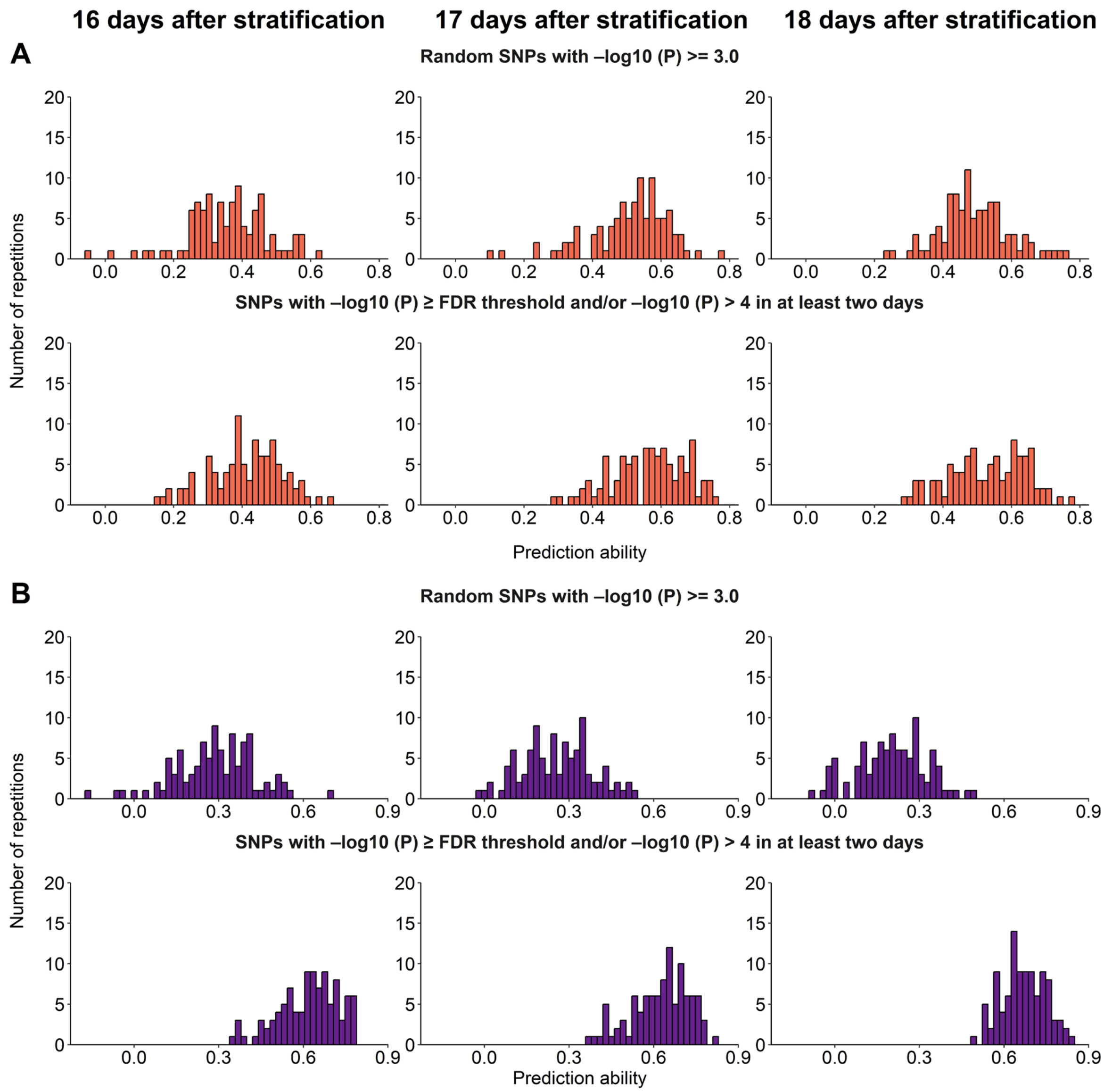


**Supplementary Fig. 5:** Distribution of prediction ability of gBLUP models for $F_q'/F_m'$ based on random GWAS-derived SNPs with -$\log_{10}$ ($P$) ≥ 3 and SNPs with -$\log_{10}$ ($P$) ≥ FDR threshold (0.05) and/or -$\log_{10}$ ($P$) > 4 on at least two days. A, 15°C $T_{night}$. B, 20°C $T_{night}$. For the plots of random SNPs with -$\log_{10}$ ($P$) ≥ 3, prediction ability was calculated by randomly selecting the same number of SNPs as for -$\log_{10}$ ($P$) ≥ FDR threshold and/or -$\log_{10}$ ($P$) > 4 on at least two days.

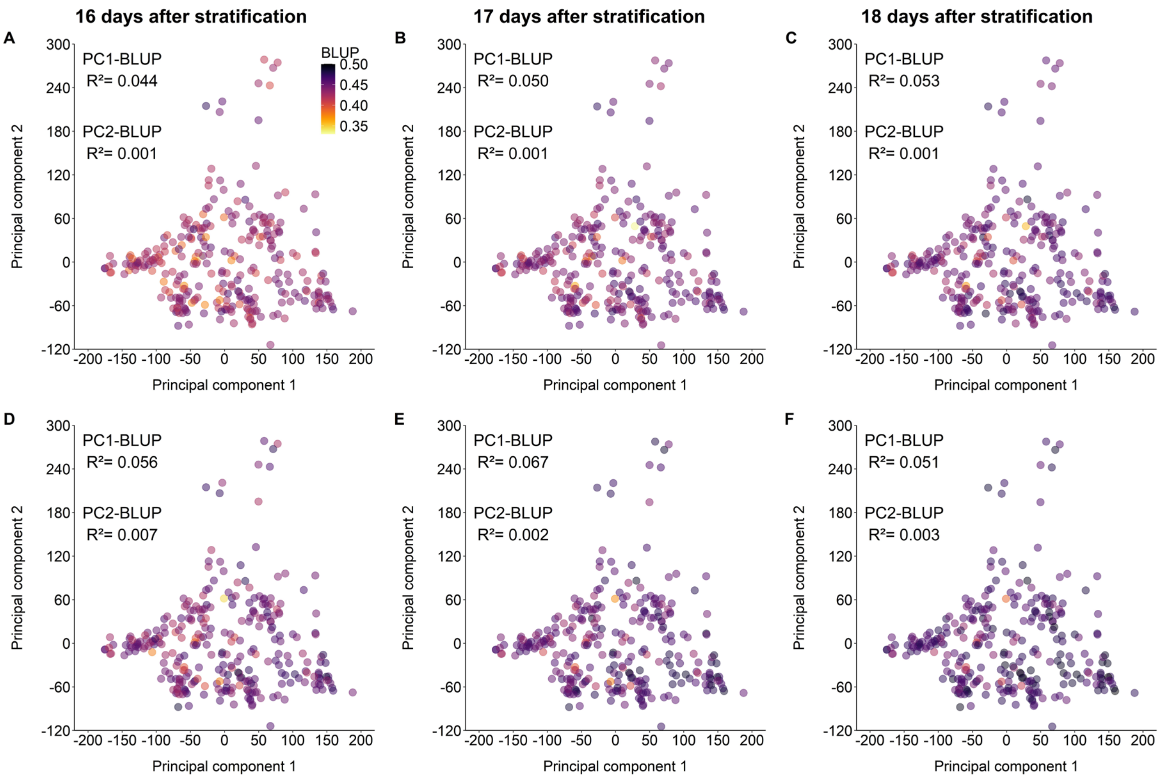


**Supplementary Fig. 6:** Population structure of the 293 accessions color-coded according to the $F_q'/F_m'$ predicted by gBLUP. A-C, 15°C $T_{night}$. D-F, 20°C $T_{night}$. Principal component (PC) analysis was based on the 211,771 SNPs. Prediction of $F_q'/F_m'$ was performed by gBLUP based on the GWAS-derived SNPs with $-\log_{10}(P) \geq 3.0$ on the respective day. The color scale in panel A shows coding scheme of the predicted $F_q'/F_m'$. $R^2$ indicates the correlation between PC1 or PC2 and the gBLUP prediction of $F_q'/F_m'$.

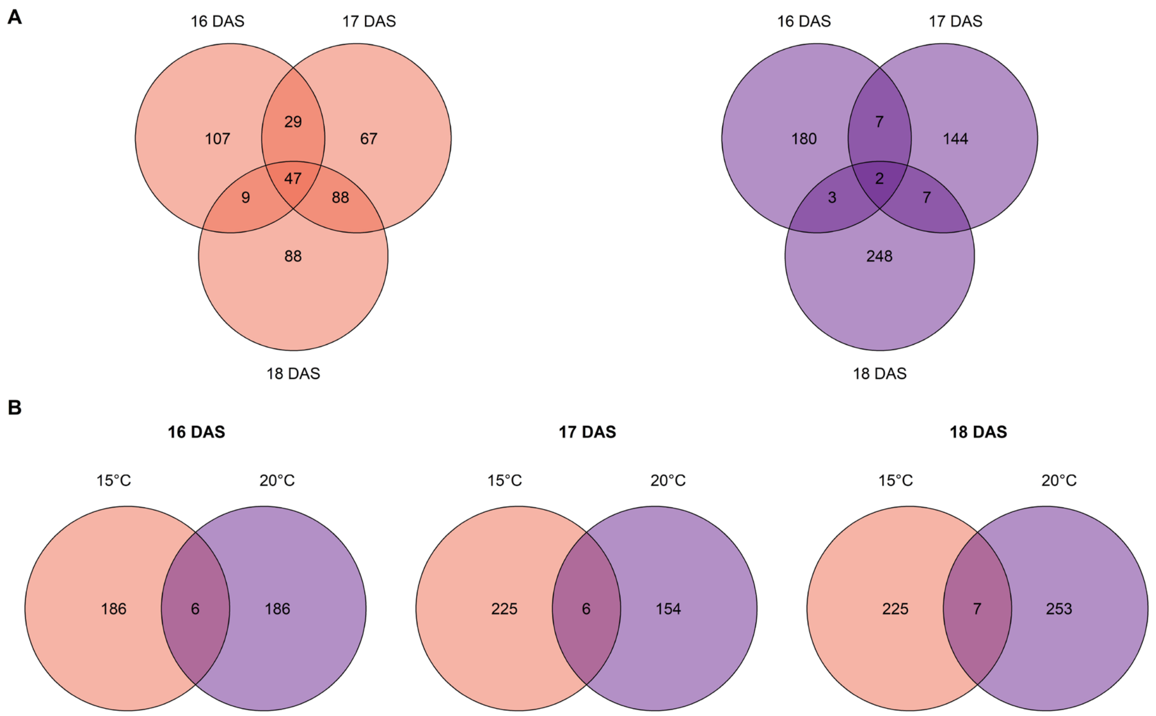


**Supplementary Fig. 7.** Shared and unique SNPs with -$\log_{10}$ (*P*) ≥ 3.0 in each $T_{night}$ condition and measurement day. Overlapping SNPs between the three measurement days for each $T_{night}$ condition (A) and between the two conditions on each day (B). Plants were growing in 15°C (coral) or 20°C (purple) $T_{night}$.

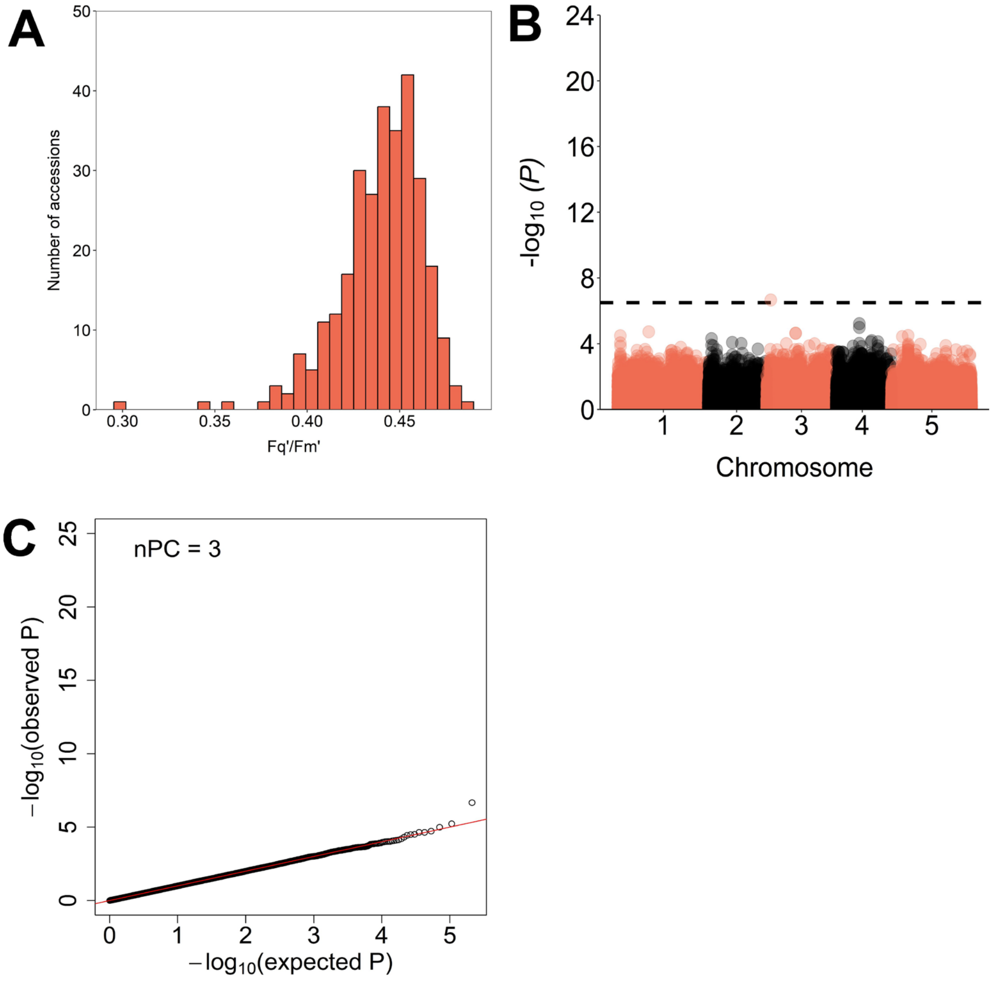


**Supplementary Fig. 8.** Analysis of $F_q$'/$F_m$' and its associated SNPs in 293 accessions in 15°C $T_{night}$ across the three measurement days combined. A, Distribution of the adjusted entry-means of $F_q$'/$F_m$' among the accessions; B, Manhattan plot; and C, Q-Q plot of the observed and the expected *P* values distribution. GWAS was performed on the adjusted entry-means of $F_q$'/$F_m$' obtained by a linear-mixed model using day as random effect. Dashed line in (B) shows the FDR (false discovery rate) threshold (0.05). nPC in (C) indicates the number of principal components included as covariates in FarmCPU.

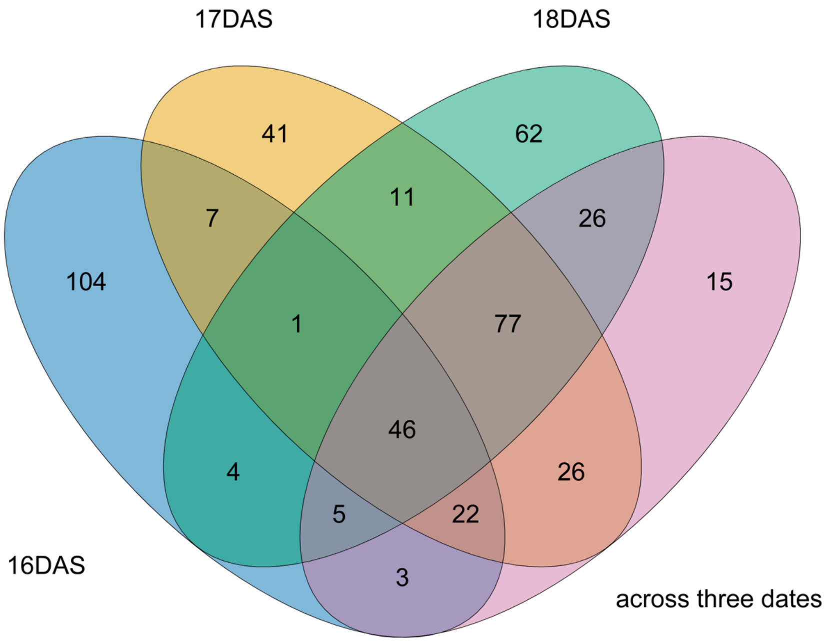


**Supplementary Fig. 9.** Shared and unique SNPs with -$\log_{10}$ ($P$) ≥ 3.0 between the three measurement days analyzed separately and across the three days combined in the 15°C $T_{night}$ condition.

.

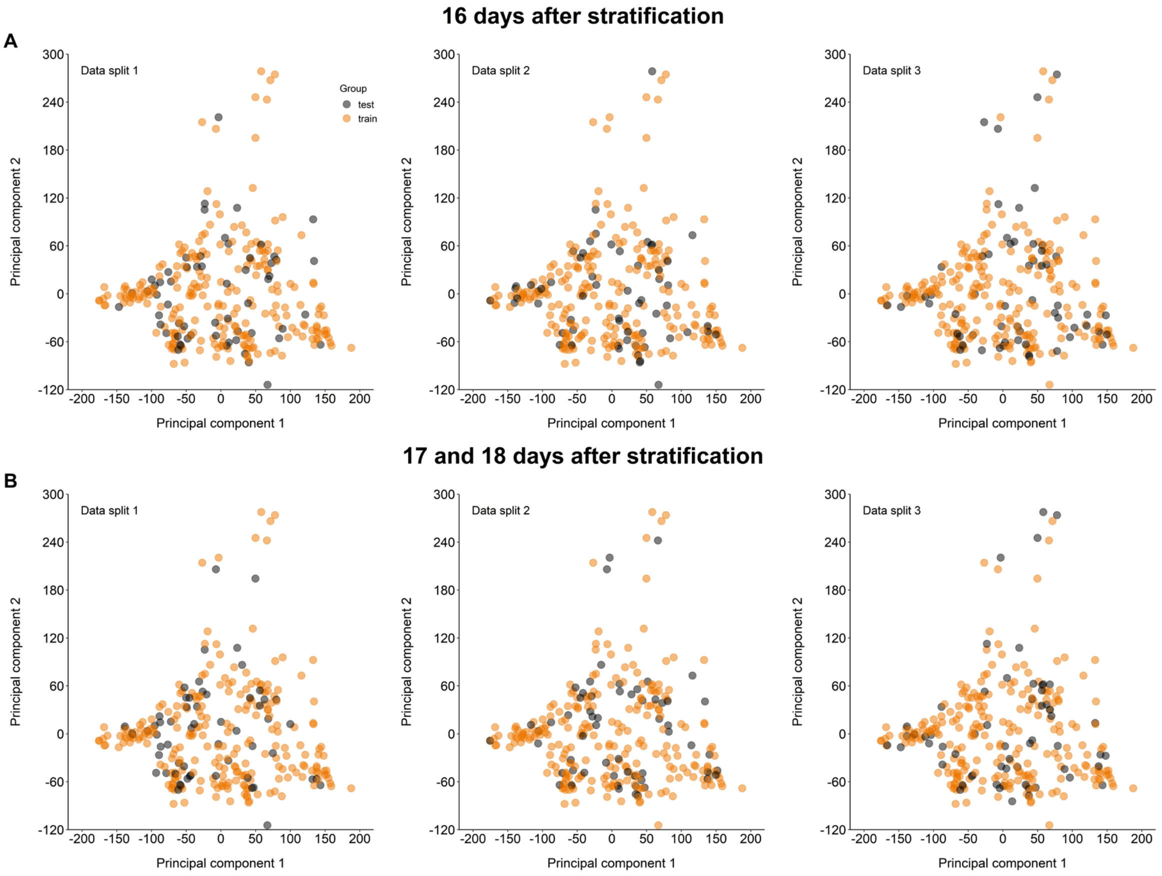


**Supplementary Fig. 10.** Population structure of randomly selected accessions for training and testing gBLUP models of $F_q'/F_m'$. The accessions were randomly split into training (orange) and test groups (black) at 4:1 ratio. The results of three test sets (three different splits) are shown for (A) 16 days after stratification (292 accessions without T1080) and (B) 17 and 18 days after stratification (all 293 accessions). Principal component analysis was based on the 211,771 SNPs.

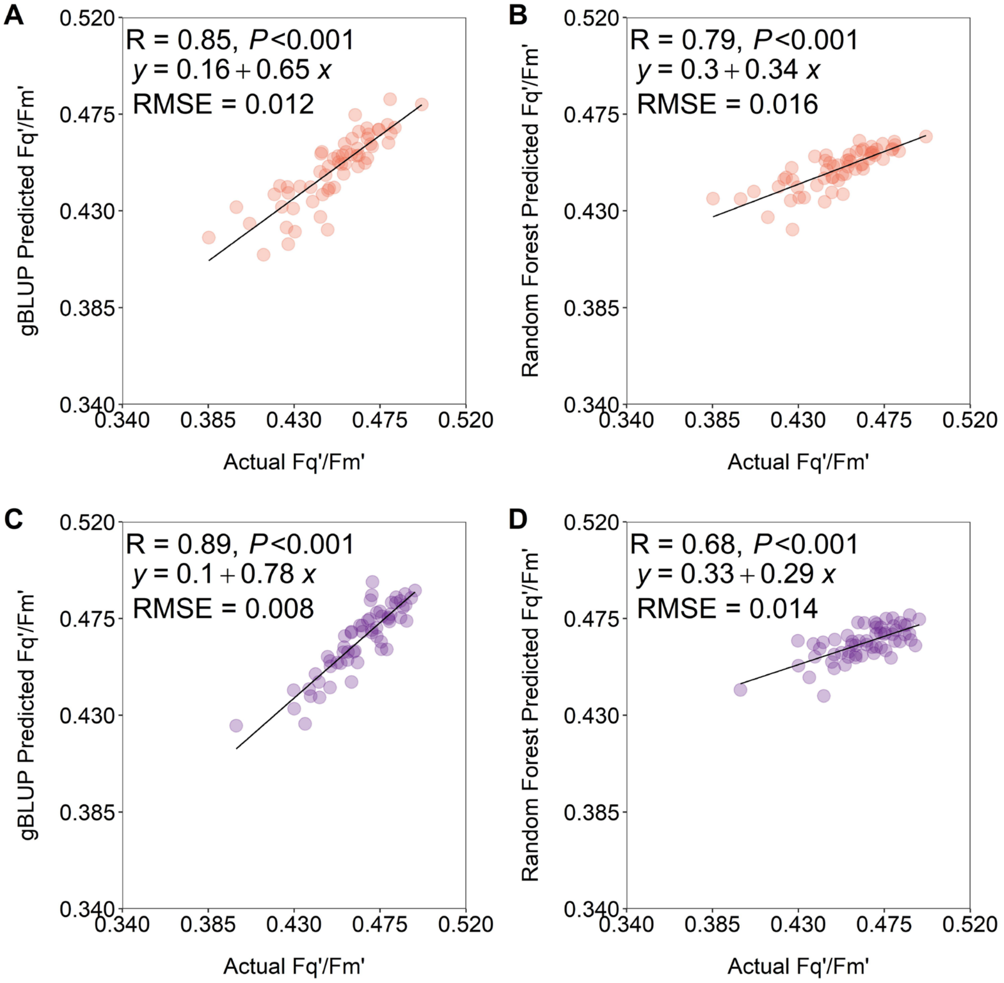


**E**

| Tnight | Test set | Method | R | RMSE |
|---|---|---|---|---|
| 15°C | 2 | gBLUP | 0.91 | 0.010 |
| | | Random forest | 0.83 | 0.017 |
| 20°C | 2 | gBLUP | 0.90 | 0.007 |
| | | Random forest | 0.66 | 0.013 |
| 15°C | 3 | gBLUP | 0.70 | 0.013 |
| | | Random forest | 0.59 | 0.014 |
| 20°C | 3 | gBLUP | 0.79 | 0.009 |
| | | Random forest | 0.61 | 0.012 |

**Supplementary Fig. 11.** Performance of gBLUP and random forest in predicting $F_q$'/$F_m$' at 18 days after stratification (DAS). The $F_q$'/$F_m$' values were predicted for the test group by gBLUP (A and C) or random forest (B and D) using the GWAS-derived SNPs with $-\log_{10}(P) \geq 3.0$ identified at 18 DAS. A and B, 15°C $T_{night}$. C and D, 20°C $T_{night}$. Circles represent 59 accessions in a test set (i.e., 1/5 of the 293 accessions randomly selected as the test group). For the other two test groups, predictive performance of gBLUP models is summarized in (E). R indicates Pearson correlation coefficient and $P$ is for significance. RMSE (root mean square error) has the same unit as $F_q$'/$F_m$'.

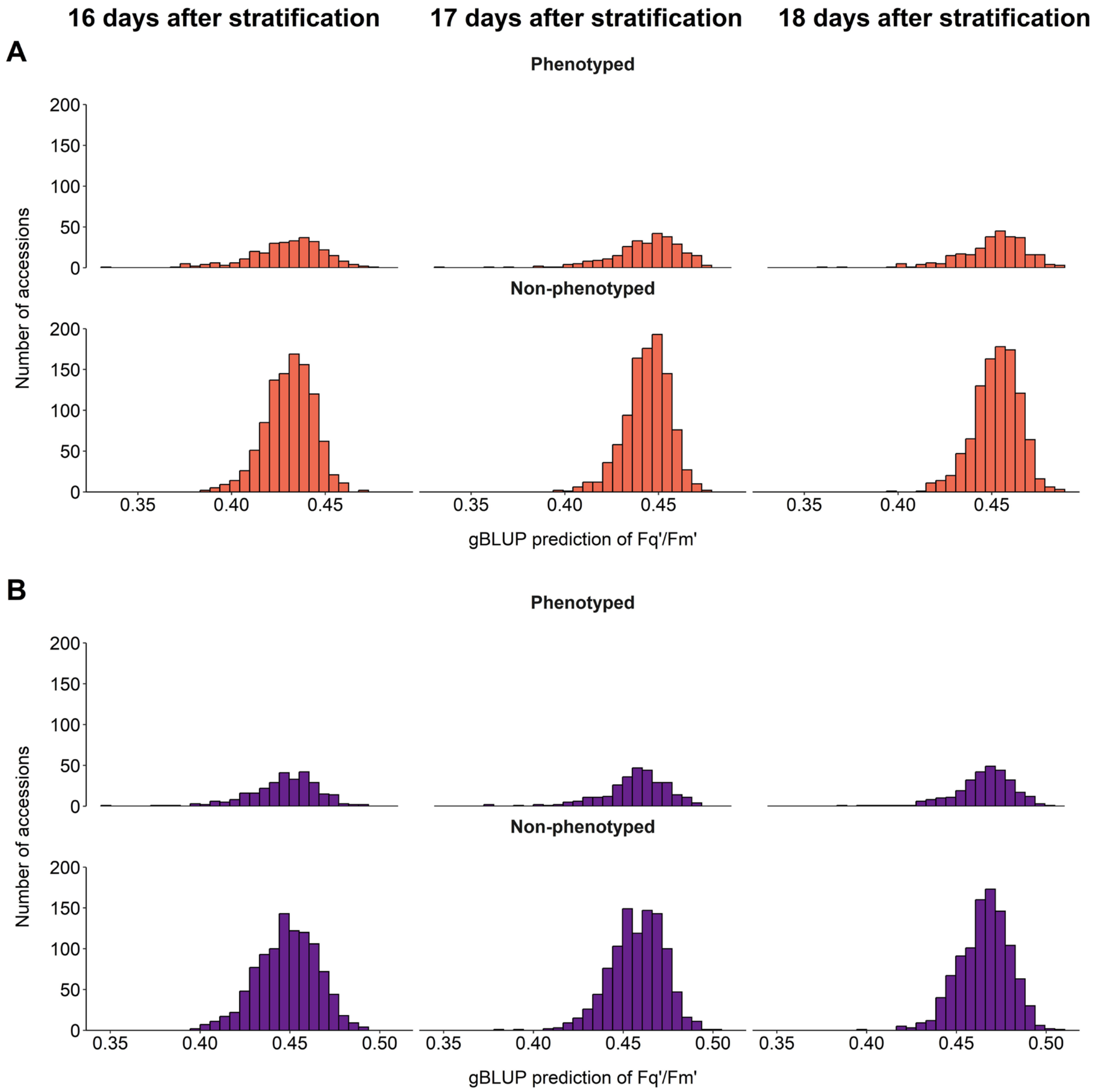


**Supplementary Fig. 12.** Distribution of predicted $F_q'/F_m'$ in the 293 phenotyped accessions in the GWAS panel and 1,014 non-phenotyped accessions from the RegMap panel. A, 15°C $T_{night}$. B, 20°C $T_{night}$. Prediction was done by gBLUP based on the GWAS-derived SNPs with $-\log_{10}(P) \geq 3.0$ identified at 16, 17 or 18 days after stratification.

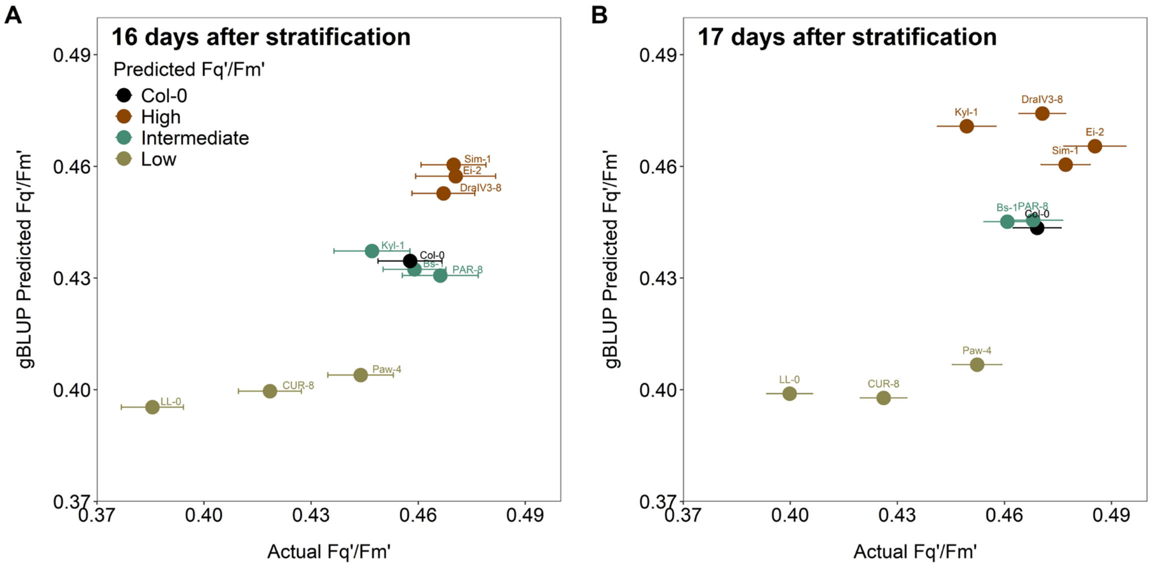


**Supplementary Fig. 13.** Experimental validation of the prediction of $F_q$'/$F_m$' by gBLUP in the 15°C $T_{night}$ condition. Predicted and actual $F_q$'/$F_m$' at 16 (A) and 17 days after stratification (B). Prediction was based on the GWAS-derived SNPs with -$\log_{10}$ (*P*) ≥ 3.0. For validation at 18 days after stratification, see Fig. 4C. Circles represent individual accessions. Different colors denote low, intermediate, and high predicted $F_q$'/$F_m$' levels. Error bars show ±SE of the adjusted entry-means of actual $F_q$'/$F_m$' of individual accessions. The number of replicate plants was: Bs-1, n = 14; CUR-8, n = 19; DralV3-8, n = 18; Ei-2, n = 3; Kyl-1, n = 4; LL-0, n = 25; PAR-8, n = 4; Paw-4, n = 11; Sim-1, n = 11; Col-0, n = 14.

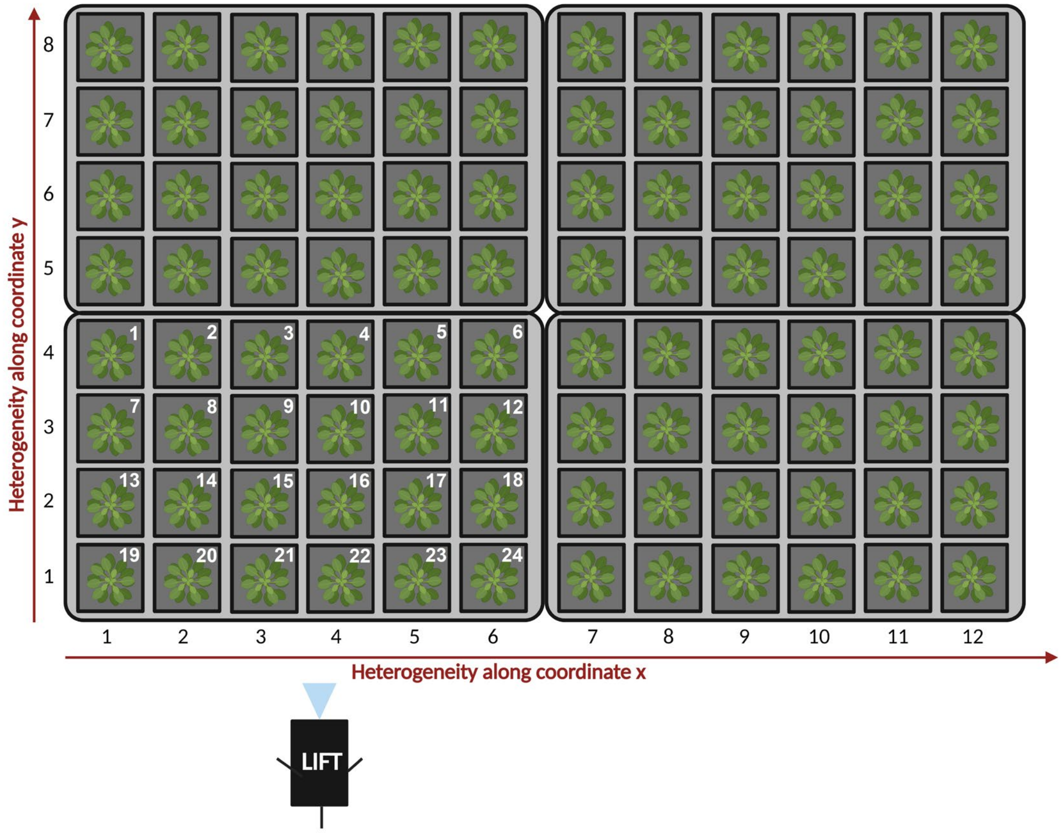


**Supplementary Fig. 14.** Scheme of plant positions in the inlets. Only four inlets are depicted as an example. To account for spatial heterogeneity of growth conditions (e.g. small differences in light intensity), two variables (coordinates x and y with numbering) were considered as random effects. For adjusting means of $F_q'/F_m'$ measured by the LIFT instrument (indicated at the left bottom), the positions of the plants inside the inlet (1-24, written in white) were considered as random effects.
